# Overcoming evolved resistance to population-suppressing homing-based gene drives

**DOI:** 10.1101/088427

**Authors:** John M. Marshall, Anna Buchman, Héctor M. Sánchez C., Omar S. Akbari

## Abstract

The use of homing-based gene drive systems to modify or suppress wild populations of a given species has been proposed as a solution to a number of significant ecological and public health related problems, including the control of mosquito-borne diseases. The recent development of a CRISPR-Cas9-based homing system for the suppression of *Anopheles gambiae*, the main African malaria vector, is encouraging for this approach; however, with current designs, the slow emergence of homing-resistant alleles is expected to result in suppressed populations rapidly rebounding, as homing-resistant alleles have a significant fitness advantage over functional, population-suppressing homing alleles. To explore this concern, we develop a mathematical model to estimate tolerable rates of homing-resistant allele generation to suppress a wild population of a given size. Our results suggest that, to achieve meaningful population suppression, tolerable rates of resistance allele generation are orders of magnitude smaller than those observed for current designs for CRISPR-Cas9-based homing systems. To remedy this, we propose a homing system architecture in which guide RNAs (gRNAs) are multiplexed, increasing the effective homing rate and decreasing the effective resistant allele generation rate. Modeling results suggest that the size of the population that can be suppressed increases exponentially with the number of multiplexed gRNAs and that, with six multiplexed gRNAs, a mosquito species could potentially be suppressed on a continental scale. We also demonstrate successful multiplexing *in vivo* in *Drosophila melanogaster* using a ribozyme-gRNA-ribozyme (RGR) approach – a strategy that could readily be adapted to engineer stable, homing-based suppression drives in relevant organisms.

## Significance Statement

Homing-based gene drive systems have the potential to rapidly invade, suppress, and eliminate wild populations of a given species. The recent engineering of a CRISPR-Cas9-based homing drive system in the main African malaria vector, *Anopheles gambiae*, highlights the potential application of these systems to global public health; however, concerns have been raised regarding the evolution of alleles resistant to the homing system, which have a significant selective advantage over functioning homing alleles. To mitigate this, we propose a design in which guide RNAs are multiplexed, reducing the emergence rate of homing-resistant alleles. Using a mathematical model, we show how this design could potentially enable population suppression on a continental scale. We also demonstrate a multiplexing design *in vivo* in *Drosophila melanogaster*.

## Introduction

The concept of using homing-based gene drive systems to rapidly invade wild populations and spread effector genes (e.g. conferring pathogen resistance) or to suppress and eliminate populations was first suggested by Burt in 2003 (1). These systems have the remarkable ability to cheat during meiosis, enabling them to rapidly spread into a population even if they confer a fitness cost to their host (2, 3). They achieve this by encoding a sequence-specific nuclease that generates a double-stranded break at one or more specific target loci in a host’s genome, directly opposite the drive. To survive, the cell is forced to rapidly repair the DNA break using its endogenous DNA repair machinery. Repair of the break using the homology-directed repair (HDR) pathway, for instance, can result in the drive system being perfectly copied into its competing allele. When this occurs in a germline cell, it effectively results in the conversion of a heterozygote into a homozygote, allowing the system to circumvent traditional Mendelian inheritance patterns and to drive into a population (2, 3). The first decade following this proposition saw moderate progress in the development of homing-based drive systems in the African malaria vector, *Anopheles gambiae* (2, 4, 5). More recently, the development of the CRISPR-Cas9 system has unlocked enormous potential for this technology, with highly functional systems being developed in quick succession to modify populations of *Drosophila melanogaster* (6), *Saccharomyces cerevisiae* (7), the Asian malaria vector, *Anopheles stephensi* (8), and the main African malaria vector, *An. gambiae* (9).

The homing-based drive systems developed using CRISPR-Cas9 have a number of highly desirable features: they are relatively straightforward to adapt to new target sequences and to port to other species, and the constructs engineered thus far have extremely high transmission rates, being inherited by 90-99% of the offspring of heterozygous parents (6–9). However, such systems are also not without their shortcomings. Firstly, at least for the CRISPR-Cas9-based constructs engineered to date, they are associated with high fitness costs, and secondly, the homing process has been shown to be highly error-prone, leading to the creation of homing resistant alleles within a few generations (8, 9). This latter shortcoming is of particular concern for population suppression strategies, because homing-resistant alleles have a strong selective advantage over functional homing alleles, leading to suppressed populations rapidly rebounding.

Homing-resistant alleles may be generated in a number of ways. For instance, they can evolve when the cell works to mend DNA damage at the target site using the non-homologous end joining (NHEJ) pathway instead of HDR following drive-induced target site cleavage. Resistant alleles may also arise due to incomplete or imperfect copying during HDR. The CRISPR-Cas9 system is particularly vulnerable to this due to its large size – the system consists of promoters, the Cas9 gene, guide RNAs and, depending on the strategy being implemented, multiple effector genes and associated regulatory elements, all of which need to perfectly copied during HDR to ensure spread into a population. Indeed, for the CRISPR-Cas9 homing construct engineered in *An. gambiae* (9), incomplete homing or internal deletion events were observed in 43% (13 out of 30) of screened organisms in which an errorless homing event was not observed. Homing-resistant alleles may also arise *de novo* via random target site mutagenesis, and some organisms may intrinsically be resistant to homing activity at a given site due to genetic variation within a species.

Therefore, while CRISPR-Cas9-based homing systems have enormous potential for the targeted engineering of populations, significant technical improvements are required if this technology is to be successfully implemented in the field (2, 10). Here, we focus specifically on the issue of homing-resistant allele generation for population suppression homing systems. We largely ignore fitness costs in this analysis as we consider these to be surmountable through tailored engineering efforts – in one of the constructs engineered thus far, fitness costs seem to result from the transgene being inserted into an eye color gene (8), and in another, due to the element copying itself to somatic as well as germline cells (9), both of which we believe to be addressable. The impact of homing-resistant alleles on homing-based population replacement strategies has been described by Noble *et al.* (11) along with a design strategy that selects against the resistance alleles. However, this solution does not apply to the population suppression systems that we explore here.

To address the impact of resistant alleles on homing-based population suppression systems, we develop a mathematical model to estimate the maximum tolerable resistant allele generation rates to achieve stable, long-term suppression for populations of various sizes. Our results suggest that, to achieve meaningful population suppression, tolerable rates of resistant allele generation are orders of magnitude lower than those observed for current CRISPR-Cas9-based homing systems. We describe how the required rates can be achieved by targeting multiple locations in a gene through guide RNA (gRNA) multiplexing (2, 3, 11, 12). Furthermore, we demonstrate successful multiplexing *in vivo* in *Drosophila melanogaster* by adapting a ribozyome-gRNA-ribozyme (RGR) approach previously demonstrated in yeast (13), and discuss possible future designs for, and challenges inherent in, the gRNA multiplexing approach for engineering stable, homing-based suppression gene drive systems. Finally, we explore the scale of population suppression that can be achieved by using this approach.

## Results

The homing population suppression system we explore here is based on that described by Hammond *et al.* (9) in which the CRISPR-Cas9 system is designed to target a gene required for female fertility. This has the effect that females homozygous for the homing allele are infertile; however, heterozygous and wild-type females have at least one functional copy of the fertility gene and hence are fertile. For sufficiently high homing rates and small fitness costs, this system is capable of spreading into a population while it reduces population fertility, eventually leading to a population crash (1). Hammond *et al.* (9) describe three strains that they engineered with this design. We consider the most successful of these – construct 7280 – for which the transmission rate from heterozygotes was ~99%, and heterozygous females had their fitness reduced by 90.7%. Approximately half (~43%) of those who did not inherit a functional homing allele from a heterozygous parent inherited a copy with errors.

### Model framework

The framework used to model this system is described in the Materials and Methods; but in short, we denote the homing allele as “H”, the wild-type allele as “h”, and the homing-resistant allele as “R”. HH females are infertile, while all other genotypes are fertile. Hh males and females produce H gametes in the germline at a frequency equal to (1+*e*)/2, where *e* denotes the efficiency of homing, or “homing rate”. Hh individuals also produce R gametes in the germline at a frequency equal to *ρ*/2, where *ρ* denotes the resistant allele generation rate. Females heterozygous for the homing allele have their fertility reduced by a fraction, *s*, while other genotypes are equally fertile (except for HH females, which are infertile). The crosses describing this system are shown in Supplementary Figure 1.

We use a discrete population, stochastic framework incorporating density-dependence at the larval stage to model this system. Our framework is modified from one previously used to examine the spread of homing endonuclease genes (HEGs) through populations of *An. gambiae* (5), the main malaria vector and the species in which the CRISPR-Cas9-based constructs were developed by Hammond *et al.* (9). This framework incorporates the egg, larval, pupal and adult life stages. Generations are overlapping and adult females mate once, retaining the genetic material of the male they mate with for the duration of their adult life. Since we are modeling a population suppression system, a discrete, stochastic model is needed to capture the chance events that happen at low population sizes. This also enables the simulation of a population crash. Density-dependence at the larval stage is important to include as it captures the phenomenon in which more larvae survive to emergence when populations are small due to reduced larval competition. To model this, we consider a monotonic increase in larval mortality with larval density.

### Expected dynamics of present constructs

With the modeling framework established, we explore the predicted dynamics of construct 7280, the best-performing construct engineered by Hammond *et al.* (9), in a population of *N* = 10,000 adult mosquitoes. The results described in Figure 1 correspond to a homing rate of *e* ≈ 98% (2 × (99% - 50%)) and a resistant allele generation rate of *ρ* ≈ 1% (~50% × (1-*e*)). The scenario in which females heterozygous for the homing allele have their fertility reduced by 90.7% is shown in Figure 1A. Here, we see that gene drive occurs slowly and population suppression is at best moderate and transient. The total adult population falls by ~42% approximately two years following a 1:1 seeding release of HH males to hh males and females; however, this suppression is short-lived – a population reduction of 30% or more is only maintained for about four months before the population rebounds.

**Figure 1.**
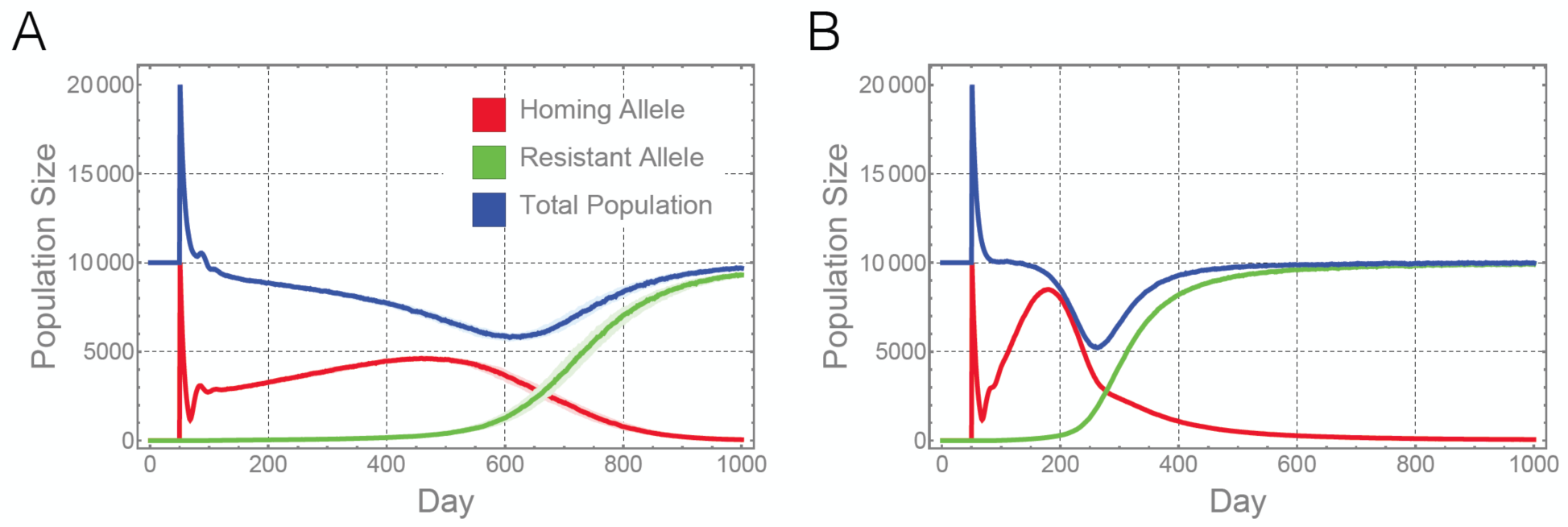
Predicted population dynamics for the present CRISPR-Cas9-based population suppression homing constructs. Here we model the predicted dynamics of the best performing construct engineered by Hammond *et al.* (9) in a population of 10,000 adult mosquitoes. The homing rate for this construct is ~98% and the resistant allele generation rate is ~1%. The model framework is described in the Materials and Methods. In panel (A), the dynamics are shown for the scenario in which females heterozygous for the homing allele have their fertility reduced by 90.7%. In panel (B), the same construct is modeled in the absence of a fertility cost. In both cases, population suppression is moderate and short-lived due to the generation of homing-resistant alleles leading to a population rebound. Red lines represent individuals having at least one copy of the homing allele (i.e. genotypes Hh, HR and HH), green lines represent individuals having at least one copy of the homing-resistant allele (i.e. genotypes hR, HR and RR), and blue lines represent the total population. Solid lines represent the median population size for 100 repetitions of the stochastic model, while shaded regions represent the 25-75% quartile range in these simulations.

Henceforth, let’s imagine that the fertility costs of the homing allele in heterozygous females can be prevented through engineering efforts to ensure that the CRISPR-Cas9 system is only expressed in germline cells. This scenario is shown in Figure 1B. Here, we see that gene drive and population suppression occur more quickly, and the extent of population suppression is slightly greater (the population is suppressed by ~48% at its peak). However, the duration of suppression is very short – a population reduction of 30% or more is only maintained for about a month. This is far less than what is hoped for gene drive-based population suppression strategies, and is a consequence of the quick emergence of homing-resistant alleles once the gene drive system becomes prevalent in the population, leading to a population rebound.

### Design criteria for population elimination

The constructs engineered by Hammond *et al.* (9) are clearly inadequate to lead to meaningful population suppression for population sizes of 10,000 adult mosquitoes; but presumably if the homing rate were increased and/or the resistant allele generation rate were decreased, then it may be possible to eliminate a specific population. In Figure 2, simulations are shown in which homing efficiency is maintained at 98% while the resistant allele generation rate is reduced from 1% (10^-2^) to 0.001% (10^-5^) to 0.00001% (10^-7^) for populations of 1,000, 10,000 and 100,000 adult mosquitoes. Here, we see that, for a resistant allele generation rate of 10^-2^ (Figures 2A-C), which is approximately what was observed for the Hammond *et al.* (9) construct, we do not expect to be able to meaningfully suppress an adult population even as small as 1,000. However, if we reduce the resistant allele generation rate by three orders of magnitude to 10^-5^ (Figures 2D-F), then we expect to eliminate populations of sizes 1,000 and 10,000, but not of size 100,000. As the resistant allele generation rate is further reduced by an additional two orders of magnitude to 10^-7^ (Figures 2G-I), we expect to eliminate adult populations of all sizes up to 100,000.

**Figure 2.**
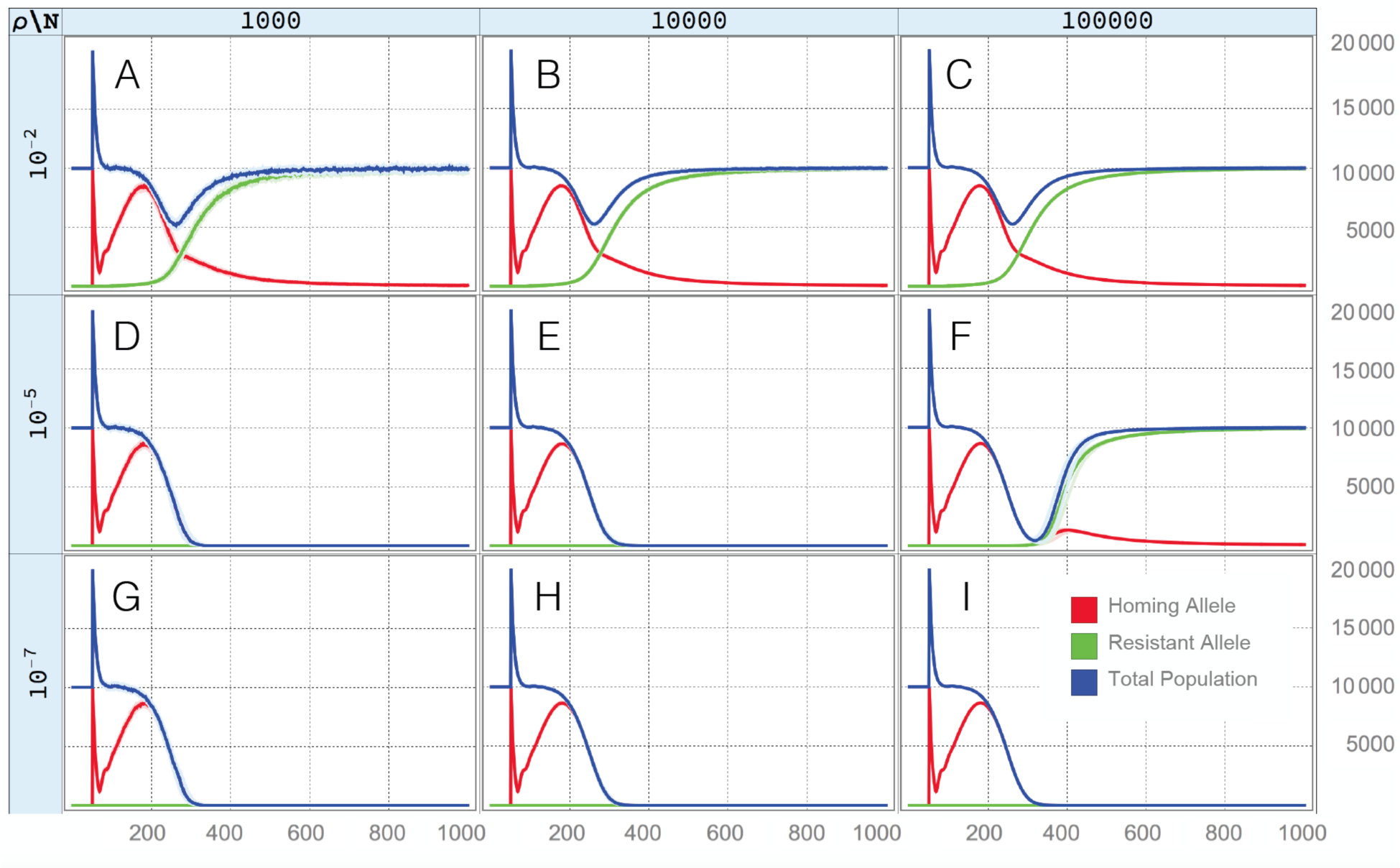
Homing and resistant allele trajectories for a range of population sizes and resistant allele generation rates. Here, we model a population suppression homing construct with a homing rate of 98% and no fertility cost. In panels (A-C) the resistant allele generation rate is 1% (10^-2^), in panels (D-F) it is 0.001% (10^-5^), and in panels (G-I) it is 0.00001% (10^-7^). In the leftmost panels (A, D and G), a population of 1,000 is modeled, in the middle panels (B, E and H), it is 10,000, and in the rightmost panels (C, F and I), it is 100,000. Red lines represent individuals having at least one copy of the homing allele, green lines represent individuals having at least one copy of the homing-resistant allele, and blue lines represent the total population. Solid lines represent the median value obtained from 100 repetitions of the stochastic model, while shaded regions represent the 25-75% quartile range. As the resistant allele generation rate is reduced, we expect to eliminate populations of larger sizes.

This trend of being able to eliminate populations of larger size with smaller resistant allele generation rates is intuitive as, in a larger population, there are more opportunities for error prone homing events to occur, leading to the emergence of homing-resistant alleles. These resistant alleles will quickly be selected for, reversing any prior population suppression. The design target for the resistant allele generation rate will therefore be determined by the population size we wish to eliminate.

The homing rate, however, does not factor into these design considerations, at least for already high homing rates. Figure 3 shows the probability of population elimination as homing efficiency is varied between 98% and 99.99% and the resistant allele generation rate is varied between 10^-2^ and 10^-7^ for a population of 10,000 adult mosquitoes. The elimination probability is calculated as the proportion of simulations (from a total of 100 per parameter set) in which the *An. gambiae* population was eliminated within 950 days of a 1:1 release of HH males to hh males and females. Here we see that, for a population size of 10,000 adults, population elimination is highly likely for resistant allele generation rates smaller than 10^-5^, and is unlikely for rates above 10^-4^. There is a critical rate between these two values at which the population is equally likely to either rebound or be eliminated and, interestingly, these dynamics are independent of the homing rate for *e* > 98%.

**Figure 3.**
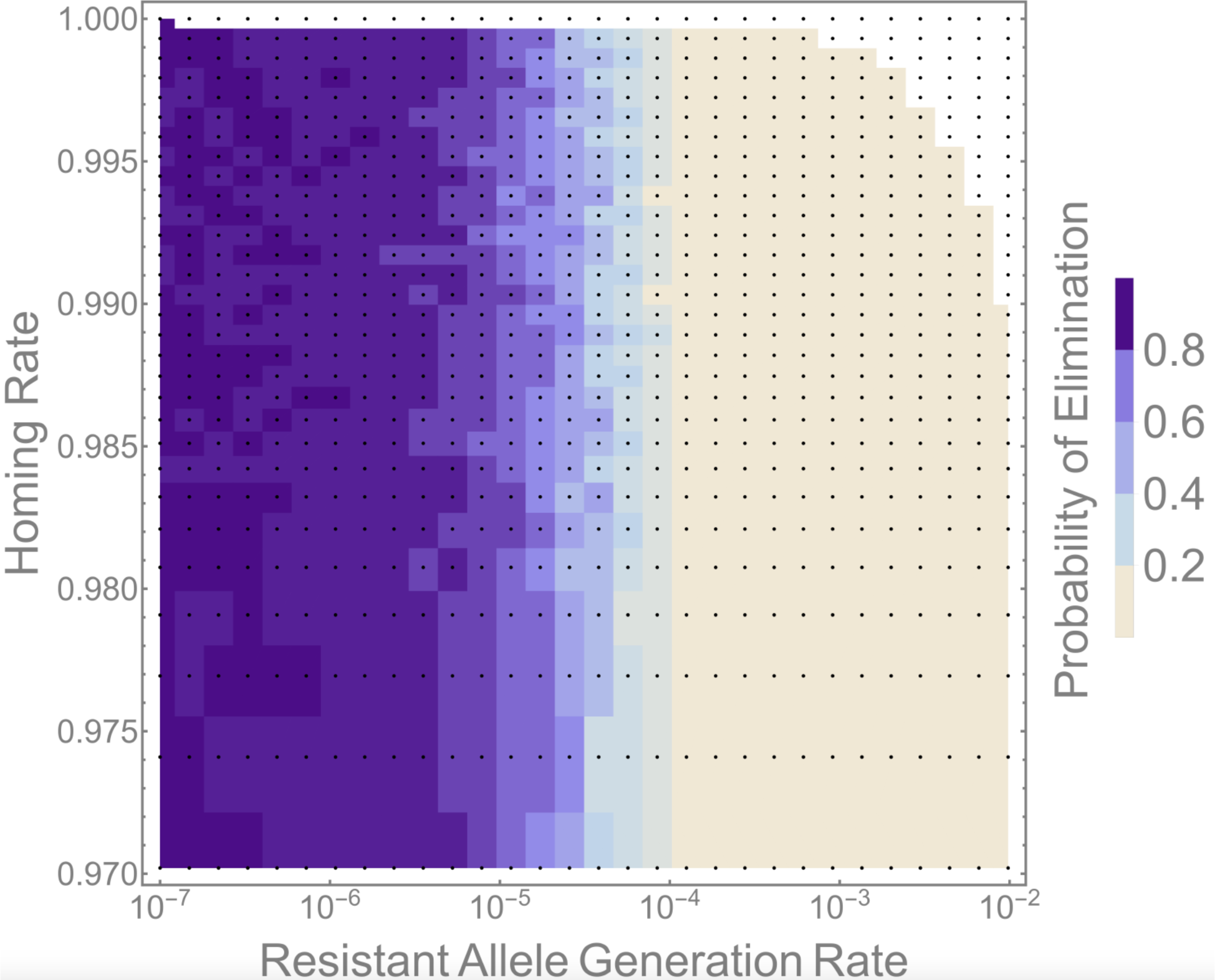
Dependence of population elimination probability on homing rate and resistant allele generation rate. Here, we model a population suppression homing construct in a population of 10,000 adult mosquitoes. Each pixel represents a combination of homing and resistant allele generation rates for which the simulation was run. Pixel shadings represent the proportion of 100 simulations in which population elimination was achieved within 950 days of a seeding 1:1 release of HH males to hh males and females. Both rate parameters were sampled logarithmically in order to gain higher resolution at high homing rates and low resistant allele generation rates. The white region represents impossible combinations of rate parameters (the rates would sum to >1). Population elimination probability is independent of the homing rate (for already high homing rates) and critically dependent on the resistant allele generation rate.

### Dependence of the design criteria for *ρ* on *N*

The independence of elimination probability and homing rate (for already high homing rates) means that we can focus our attention on achieving a resistant allele generation rate, *ρ*, small enough such that population elimination is likely for a given population size, *N*. Stochastic simulations become highly computationally intensive as population size increases, and so we seek a relationship between *N* and the corresponding resistant allele generation rate, *ρ*, for which we can be 90% sure of achieving population elimination (or sure with some other degree of certainty). To this end, Figure 4A depicts elimination probability as a function of *ρ* as we vary *N* between 1,000 and 100,000. The familiar case of a population size of 10,000 is shown in light gold and indicates that elimination is ~10% likely for a *ρ* value of 10^-4^ and ~80% likely for a *ρ* value of 10^-5^. As the population size increases from 1,000 to 100,000, we see that the *ρ* value required to achieve an elimination probability of 90% or higher becomes significantly smaller.

**Figure 4.**
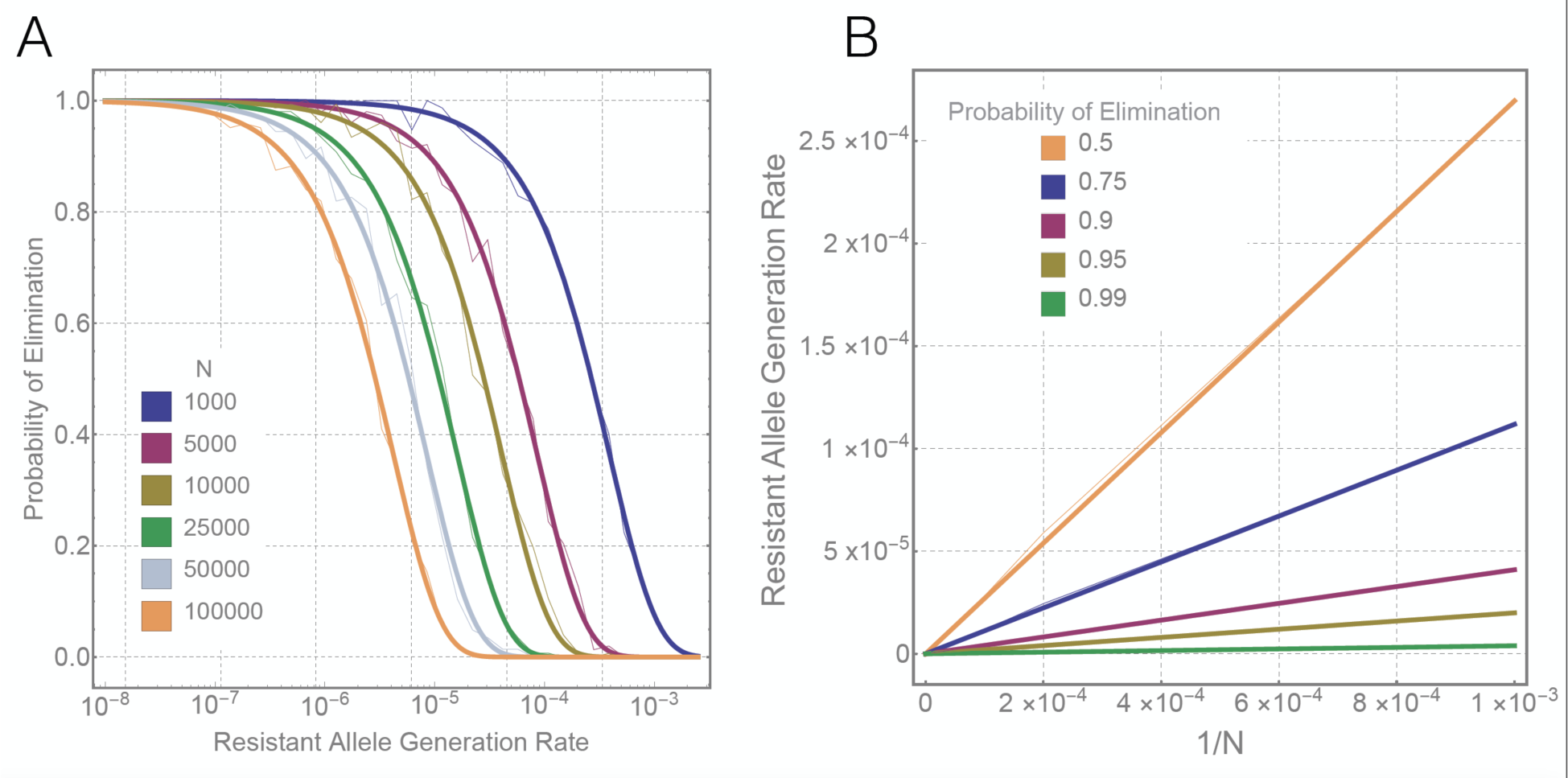
Relationship between population size and the resistant allele generation rate required for a given population elimination probability. (A) Elimination probability as a function of resistant allele generation rate for a range of population sizes*, N*, between 1,000 and 100,000. Sigmoidal curves are fitted to data points covering 30 resistant allele generation rates sampled logarithmically between 10^-2^ and 10^-7^. (B) Linear relationship between 1/*N* and the resistant allele generation rate leading to a given probability of population elimination. Values of 1/*N* are as shown in panel A, and resistant allele generation rates are inferred from the sigmoid curves. Faint lines in both panels represent interpolation between simulated data points while solid lines represent fitted linear relationships. There is a clear linear relationship between 1/*N* and the resistant allele generation rate leading to a given elimination probability.

The form of the relationship between *N* and *ρ*_*x*_, the resistant allele generation rate leading to an elimination probability of *x*, is depicted in Figure 4B for selected elimination probabilities. Fortunately for our ability to extrapolate to larger population sizes, there is a linear relationship between 1/*N* and *ρ*_*x*_ for all elimination probabilities investigated. This is understandable since each mosquito presents an opportunity for a resistant allele to emerge and prevent population elimination. Given this relationship, to be 90% sure of population elimination in a population of size *N*, the design criteria for the resistant allele generation rate is:

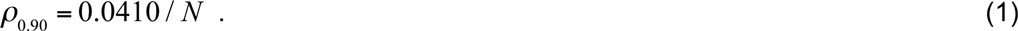

To be 95% sure of eliminating a population of size *N*, the design criteria is:

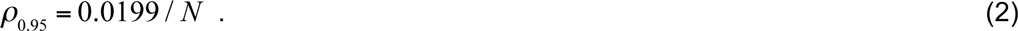

And to be 99% sure of eliminating a population of size *N*, the criteria is:

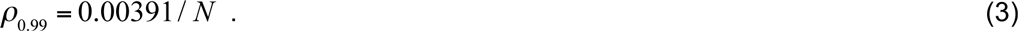

This means that, to be 90% sure of eliminating a population of size 1,000, *ρ* should be less than 4.1 × 10^-5^ (0.0041%), and to be 90% sure of eliminating a population of size 100,000, *ρ* should be less than 4.1 × 10^-7^ (0.000041%). To be 95% sure of eliminating populations of these sizes, *ρ* should be about half that predicted for a 90% chance of elimination, and for a 99% chance of elimination, *ρ* should be an order of magnitude smaller than that predicted for a 90% chance of elimination.

The population size that we wish to eliminate will vary depending on our goals; but one proposition for homing-based gene drive has been to eliminate a disease vector species such as *An. gambiae* on a continental scale. Assuming there are about ten times as many *An. gambiae* mosquitoes on the African continent as there are people, this suggests a population of ~10 billion (10^10^). In order to be 90% sure that resistant alleles will not interfere with eliminating a population this size, *ρ* should be less than 4.1 × 10^-12^ (0.0000000004%). To be 95% sure, *ρ* should be less than 2.0 × 10^-12^, and to be 99% sure, *ρ* should be less than 3.9 × 10^-13^.

### Multiplexing gRNAs

The *ρ* values required to prevent resistant alleles from interfering with the elimination of an *An. gambiae* population on the scale of the African continent are vanishingly small; but interestingly, the *ρ* value required to have a 90% chance of suppressing a population of just 1,000 adult mosquitoes is already several orders of magnitude smaller than that observed for the best performing construct of Hammond *et al.* (9). Given the inevitability of the evolution of homing resistant alleles, mitigating their impact is imperative to creating functional and stable homing based population suppression gene drive systems.

A promising strategy for achieving this feat is the multiplexing of gRNAs in the gene drive to target multiple sequences. This idea has been previously proposed to increase the stability of drives (2, 3, 11, 14); however, to date only one multiplexing strategy has been demonstrated to function in a whole-animal model (12). Therefore, to further expand the toolbox for multiplexing gRNAs in whole animals, here we test effectiveness of a technique previously demonstrated in yeast that relies on flanking gRNAs with self-cleaving ribozymes, known as the ribozyme-gRNA-ribozyme (RGR) approach *in vivo* in *Drosophila melanogaster* (13). To validate this technique, we generated plasmid OA-16 that contains two multiplexed RGRs, the first targeting the *white* gene and the second targeting the *yellow* gene at target sequences previously validated (15), driven by a single polymerase-2 ubiquitin promoter (16) (Figure 5A-B). The plasmid also contains a *white* gene that could be targeted by the *white* gRNA and used for detecting transgenic individuals bearing the OA-16 construct.

**Figure 5.**
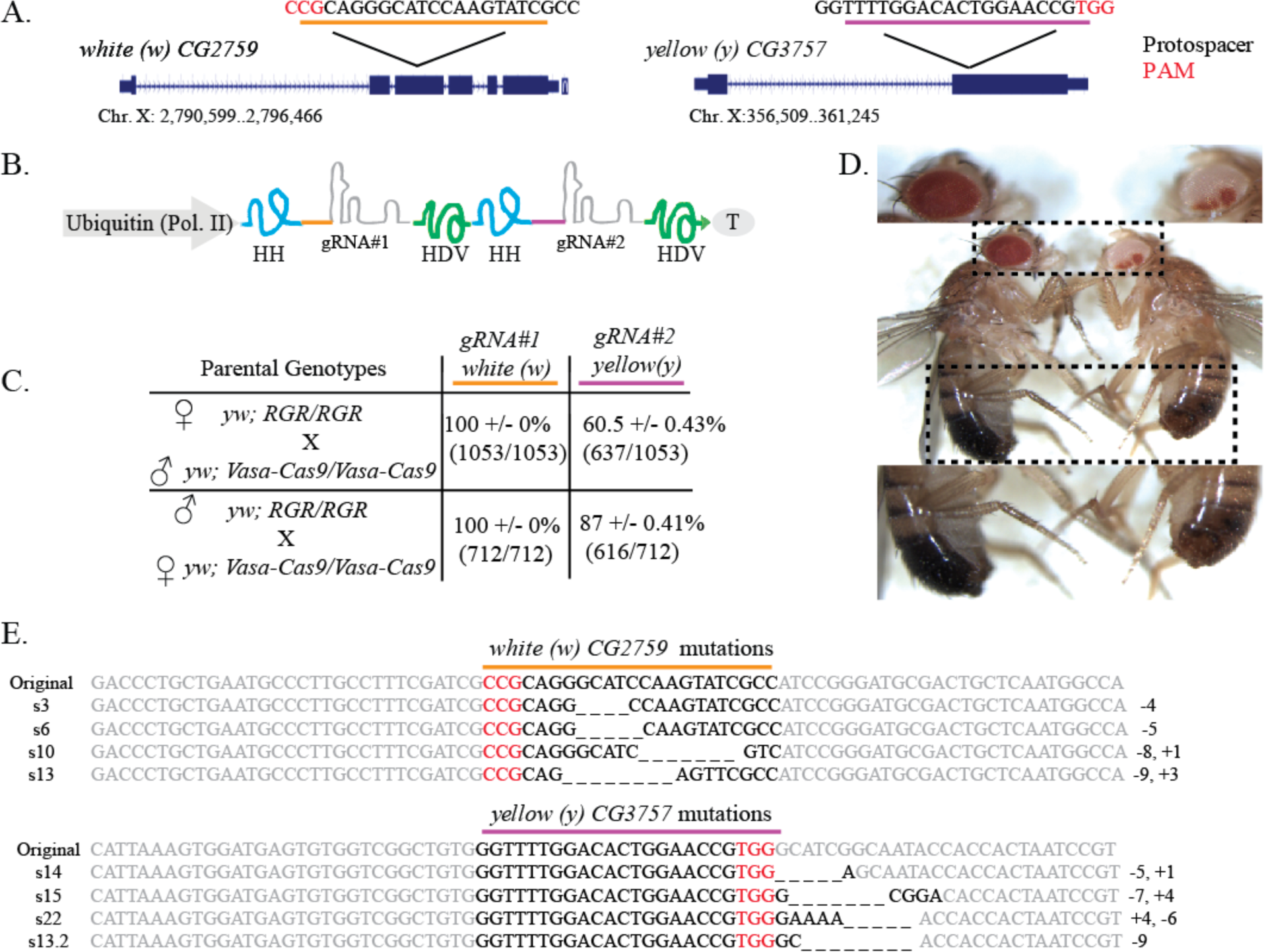
RGR/Cas9-induced mutations at the yellow and white loci. (A) Schematic of the white and yellow genes showing the gRNA target sites. Exons are shown as blue boxes, the gRNA target site locations are indicated by black lines, and the gRNA target site sequences (with black letters indicating protospacer sequences and red letters indicating PAM sequences) are underlined in yellow for white, purple for yellow. (B) Schematic of the OA-16 construct utilized in generating mutations. The first and second gRNAs (targeting white and yellow, respectively) are shown in grey. Each gRNA has a hammerhead ribozyme 5’ (shown in blue) and an HDV ribozyme 3’ (shown in green). The gRNAs are driven by a single *Drosophila* ubiquitin polII promoter. (C) Crossing scheme used to generate mutants, and obtained results. Individual male and female flies homozygous for the OA-16 construct were crossed to individual female and male flies, respectively, of a homozygous vasa-Cas9 line. Progeny were scored for eye and body color. Percentages correspond to number of flies out of total cross progeny (+/- SEM) exhibiting a mutation for each gRNA. (D) The white and yellow tissues of an OA-16 homozygous male fly with no exposure to Cas9 are un-mutated (left), while the white and yellow tissues of a fly generated by crossing OA-16 homozygotes to vasa-Cas9 homozygotes show mosaic expression (right). (E) Examples of sequences of CRISPR/Cas-induced mutations in white (top) and yellow (bottom). The first line in each alignment represents wild-type sequence, and subsequent lines show individual mutant clones.

This plasmid was integrated site-specifically into the *Drosophila attP* line BSC (Bloomington Stock Center) 24486. Generated transgenic males and females were individually mated to females and males of transgenic line BSC 51324 (*vasa-Cas9*), and the progeny of the resulting crosses were scored (Figure 5C). As expected, transformant flies bearing the OA-16 plasmid had no mutations in the *white* or *yellow* genes in the absence of Cas9 (Figure 5D, left). All of the scored offspring (712 from OA-16 male/Cas9 female crosses and 1053 from OA-16 female/Cas9 male crosses) had white or variegated eyes (Figure 5D), indicating that the first of the two RGRs had a cleavage efficiency near 100%. Additionally, 87% +/- 0.41 (616/712) of the offspring of OA-16 male/Cas9 female crosses and 60.5% +/- 0.43 (637/1053) of the offspring from OA-16 female/Cas9 male crosses had a predominantly yellow cuticle (Figure 5D, right), indicating that the second RGR was also functional, albeit with a significantly lower cleavage efficiency than the first. The presence of mutations was confirmed by sequencing of PCR products that span the cleavage site (Figure 5E). Together, these data conclusively provide a proof-of-principle for the feasibility of the RGR approach as a method for multiplexing gRNAs in whole animals.

### Design requirements for multiplex number

This demonstration of the multiplexing of gRNAs is encouraging for the same being achieved in insect species that transmit human diseases, such as *An. gambiae*. Multiplexing is expected to increase the effective homing rate, as only one of several target sites must have a functional copy of the homing allele in order for the composite allele to have the homing phenotype. That said, as depicted in Figure 3, the probability of population elimination is independent of the homing rate for *e* > 98%, but is highly dependent on the resistant allele generation rate, *ρ*. We derive the effective resistant allele generation rate for two and three multiplexed gRNAs in Supplementary Text S1 and find that, for a multiplex number of *m*, this is approximately equal to *ρ_m_* in both cases. This logically follows since resistant allele generation in the presence of multiplexing requires all gRNA target sites to have a homing-resistant allele. In Supplementary Text S1 we show that, although homing-resistant alleles may accumulate in a composite allele with multiple target sites, partially resistant composite alleles are rarely generated and are frequently converted to homing alleles soon after they have been formed. The rate of completely resistant composite alleles emerging is therefore approximately equal to the rate of resistant alleles emerging at all target sites at once, i.e. *ρ_m_*.

The effective resistant allele generation rate therefore becomes exponentially smaller as the number of multiplexed gRNAs increases. For a baseline *ρ* value of 1%, the effective *ρ* value becomes ~10^-4^ for a multiplex number of two, ~10^-6^ for a multiplex number of three, and ~10^-2^*m* for a multiplex number of *m*. Following on from Equation 1, for a baseline resistant allele generation rate, *ρ*, the multiplex number, *m*, we must achieve in order to have a 90% chance of eliminating a population of size *N* is given by:

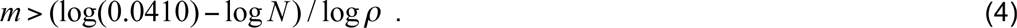

To be 95% sure of eliminating a population of size *N*, the criteria is:

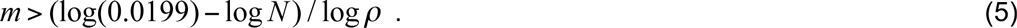

And to be 99% sure of eliminating a population of size *N*, the criteria is:

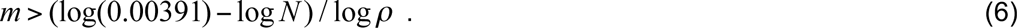

This means that, to have a 90% chance of eliminating a population of size 10,000, we require a multiplex number of three; for a population of size 1 million, we require a multiplex number of four; and for a population of size 10 billion, we require a multiplex number of six. An encouraging property of these predictions is that, as multiplex number increases linearly, the population size that we can eliminate increases exponentially. A modest additional increase in multiplex number can also lead to a much higher chance of eliminating a population of the same size. For instance, a multiplex number of seven is predicted to provide a >99% chance of eliminating an *An. gambiae* population on the scale of the African continent (~10 billion). Important spatial factors are not considered here.

In the multiplexing experiments demonstrating the RGR approach in *D. melanogaster*, a reduced cleavage rate was observed at the second target site as compared to the first. Presumably this is something that could be solved through subsequent engineering efforts; however, mathematical analysis described in Supplementary Text S1 shows that, at least for the two-gRNA system, a reduced cleavage rate at one site doesn’t significantly alter the effective resistant allele generation rate overall. This is because a reduced cleavage rate at one site also suggests a reduced resistant allele generation rate at this site, as both H and R alleles are generated through the same cleavage and repair mechanism. However, this reduction in the resistant allele generation rate is compensated for by the increased accumulation of partially resistant composite alleles and their subsequent development into completely resistant composite alleles. The net effect is that, even if one of two multiplexed gRNAs displays a reduced cleavage rate, the benefits of multiplexing in terms of reduced resistant allele generation are very similar.

## Discussion

The possibility of using gene drive systems to suppress and potentially eliminate wild populations has provoked intense interest over the last decade (2, 3, 17–19). This excitement has recently been fueled by significant developments in genetic engineering, and in particular by the CRISPR revolution, which has enabled scientists to develop homing-based gene drive systems targeting a range of sites in any genome with relative ease. In terms of population suppression, recessive lethal and sterility genes are of particular interest as targets because a homing system targeting these genes can potentially spread to fixation and eliminate the population in the process, even when introduced beginning with a single drive-containing organism (4). While this excitement is warranted, it is highly relevant to determine how the evolution of homing-resistant alleles could interfere with population suppression drive strategies, and to determine design criteria to increase stability of the drive for the likely elimination of populations of a given size.

To address this important question, we developed a mathematical model to describe the spread of a population-suppressing homing allele through a population of *An. gambiae*, and the impact that a homing-resistant allele could have on these dynamics. Homing-resistant alleles may originate through several mechanisms: a) *de novo* mutations, which occur independently from the drive; b) pre-existing natural variation in the population, which may be minimized through intelligent selection of homing recognition sites; and c) in response to the drive, by the cell’s utilization of the endogenous DNA breakage repair machinery to mend DNA damage caused by the drive via non-homologous end joining (NHEJ). While the former two mechanisms are important, we have focused this study on the latter (i.e. resistant allele formation in response to the drive), as this is expected to occur at a significantly higher frequency than the other mechanisms (20).

We discover that, despite promising experimental data reporting extremely high rates of homing in the germline for recently engineered CRISPR-Cas9-based homing systems (6, 8, 9), population suppression will be at best moderate and short-lived with current construct architectures due to the quick generation of homing-resistant alleles. Our predictions are that, to prevent a population rebound, presently observed homing rates are adequate; however, reducing the resistant allele generation rate is critical. For example, to be 95% sure that resistant alleles will not interfere with suppression of a population of *An. gambiae* mosquitoes on the scale of the African continent, the resistant allele generation rate should be less than ~2 × 10^-12^ per homing event – about 10 orders of magnitude smaller than presently-observed resistant allele generation rates (9).

While it might be near impossible to achieve resistant allele generation rates this low with a single gRNA recognizing an exclusive target site, one strategy to mitigate the impact of homing resistant alleles is to multiplex gRNAs in the drive system (2, 3, 9, 14). By multiplexing gRNAs to target multiple locations within an essential gene, each site is required to be homing-resistant in order for the composite allele to have the homing-resistant phenotype. Our results suggest that the effective resistant allele generation rate becomes exponentially smaller as the number of multiplexed gRNAs increases, and that a multiplex number of six may be sufficient to have a 90% chance of eliminating an *An. gambiae* population on the scale of the African continent.

Several approaches to multiplexing gRNAs have been described, including the use of different polIII promoters such as U6:1-U6:3 (21), HP1 (22, 23), 7SK (22), or tRNA promoters (24) to promote expression of individual gRNAs (Figure 6A-B). While these strategies are effective, they are limited by the fact that most polIII promoters do not drive temporal and/or tissue specific expression, which may incur increased fitness costs to the organism due to ubiquitous and continuous gRNA expression. These strategies also require an individual promoter element for each gRNA, thereby increasing the overall size of the drive and possibly introducing repetitive elements. Repetitive DNA sequences have reduced stability (25) and previous attempts to build drives with zinc-finger nucleases and TALENs have indeed demonstrated that larger and more repetitive the drive systems are less evolutionarily stable. Therefore, it is essential to minimize the size and repetitiveness of the drive (26). To circumvent the need to express each gRNA from a different polIII promoter, gRNAs can be flanked with self-cleaving ribozymes (13, 24, 27, 28) or tRNAs (12, 21, 24, 29), which can allow the use of a single temporal and tissue-specific polII promoter to drive expression of an array of flaked gRNAs, thus reducing the overall drive size and repetitiveness of the drive element.

**Figure 6.**
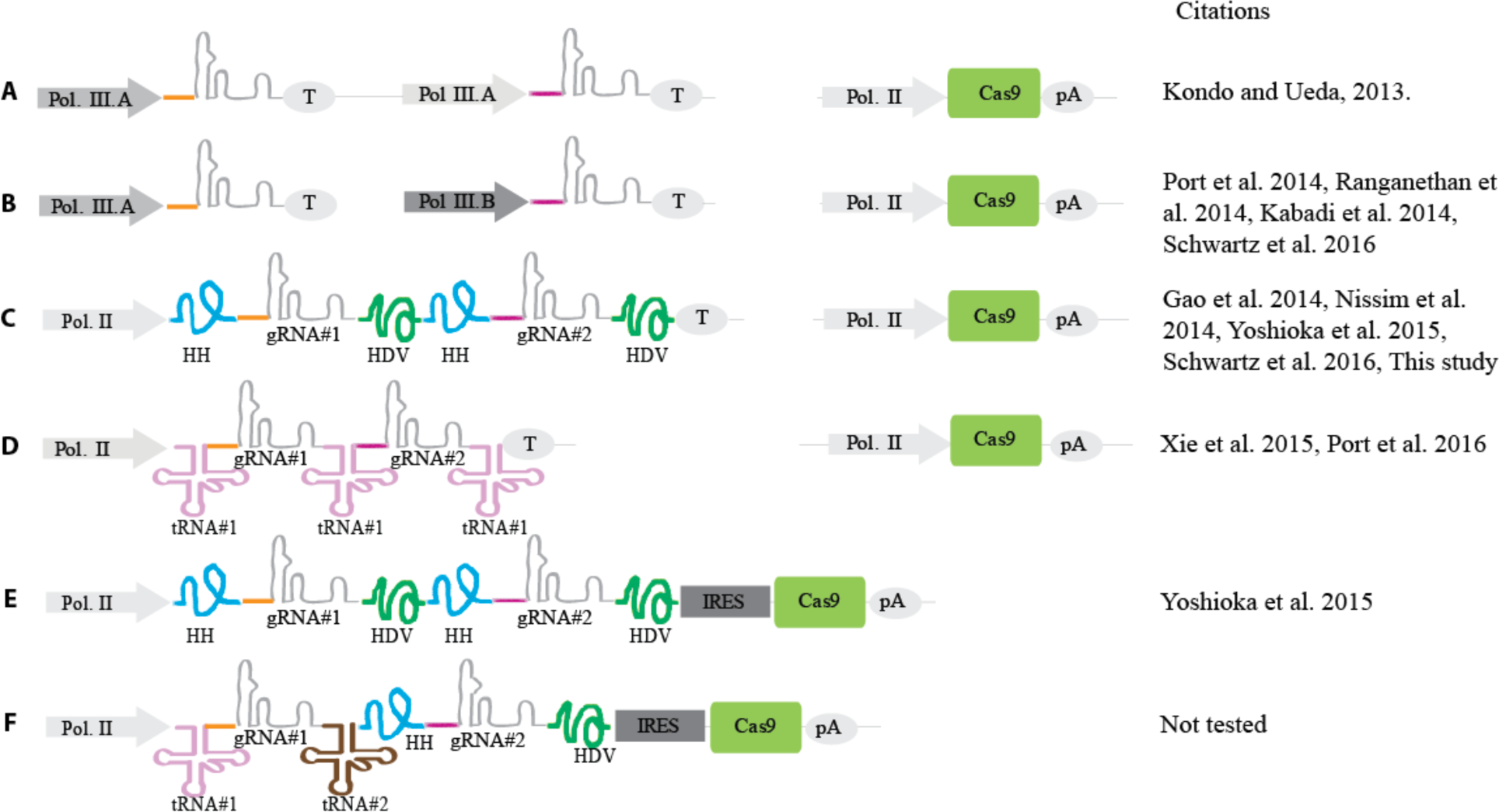
Schematic of various proposed strategies for multiplexing gRNAs. (A) A gRNA multiplexing scheme where the same polIII promoter drives each of two gRNAs, and a polII driven Cas9 is provided as a separate transgene. (B) A multiplexing scheme where two different polIII promoter drive each of two gRNAs, and a polII-driven Cas9 is provided as a separate transgene. (C) A multiplexing scheme where each of two gRNAs are surrounded by a 5’ HH ribozyme and a 3’ HDV ribozyme, and a polII-driven Cas9 is provided as a separate transgene. (D) A multiplexing scheme where the two gRNAs are surrounded by copies of the same tRNA (with a tRNA 5’ of the first gRNA, between gRNAs 1 and 2, and 3’ of the second gRNA), and a polII-driven Cas9 is provided as a separate transgene. (E) A multiplexing scheme where each of two gRNAs are surrounded by a 5’ HH ribozyme and a 3’ HDV ribozyme, as in (C), but the Cas9 is located on the same transgene, 3’ of the gRNAs and preceded by an IRES. (F) A proposed multiplexing scheme where the first of two gRNAs is surrounded by two different tRNAs, the second gRNA is flanked by the HH and HDV ribozymes (as in (C) and (E)), and the Cas9 is located on the same transgene, 3’ of the gRNAs and preceded by an IRES (as in (E)). Grey triangles represent polIII promoters; blue triangles are polII promoters; terminators (T) and polyA signals (pA) are shown in grey ovals; Cas9 is represented as a green rectangle; internal ribosomal entry sequences (IRES) are grey sequences; gRNA scaffolds are shown as grey lines, with red and purple connecting lines representing two different gRNAs; the hammerhead (HH) and HDV ribozymes are shown as blue and green lines, respectively; and two different tRNAs (tRNA^Gly^ and a non-specific tRNA) are shown as pink and brown lines, respectively.

To date, only the tRNA gRNA multiplex approach has been validated in a whole animal model (12). Here, we report that the adaptation of another multiplex RGR approach, previously demonstrated in yeast (13), functions efficiently in the *D. melanogaster.* We observe highly efficient cleavage rates approaching 100% for the first gRNA and of 60%-86% for the second gRNA, depending on whether Cas9 is either maternally or paternally inherited (Figure 5). Importantly, while we use the same two ribozymes to flank each gRNA (Figures 5B & 6C), the RGR approach may be expanded in the future to incorporate different ribozymes than the two tested here (30, 31) to reduce repetitiveness, and the same strategy could be applied to the use of tRNAs other than the tRNA^Gly^ (Figure 6D) that was shown to work by Port *et al.* (12). Furthermore, as recently demonstrated by Yoshiokda *et al.* (28) in mammalian cells, the RGR approach could also be optimized to allow for expression of both the multiplexed gRNAs and CRISPR-Cas9 components from a single polII promoter, further reducing drive size and repetitiveness (Figure 6E). Utilization of this single promoter-gRNA-CRISPR/Cas9 strategy combined with both the tRNA and RGR approaches may yield an optimal drive element design, both in terms of efficiency and stability (Figure 6F). Finally, it may be important to utilize improved gRNA backbones to further reduce repetitiveness of the gRNAs (32). Overall, the above design considerations may offer opportunities to engineer compact, evolutionarily stable gene drive cassettes; however, these ideas are largely untested and the use of multiplexed gRNAs in a functional gene drive system remains to be demonstrated.

Several assumptions have been made in the modeling portion of this study. Most noteworthy is the description of an *An. gambiae* population on the scale of the African continent as randomly mixing. Clearly, the study of gene drive in *An. gambiae* at anything beyond the village scale will require an understanding of population structure, and in fact, even at the village scale there are population considerations regarding gene flow within the *An. gambiae* species complex (33, 34). By ignoring population structure, the model described here cannot be used to make claims regarding the time course or spatial pattern of gene drive on a large scale (35). However, despite this, each target site on a chromosome represents an opportunity for homing resistance to emerge and, in this sense, we expect there to be some validity to predictions regarding the resistant allele generation rate required to make a population rebound unlikely.

At the molecular level, several assumptions have been made regarding the dynamics of multiplexed gRNAs. In particular, these have been modeled as independently-acting homing systems; however, the use of multiplexed gRNAs in a functional gene drive system has yet to be demonstrated and hence these dynamics will be elucidated in future drive experiments. Potential problems may arise from sequence repetitiveness in the drive element if identical gRNA backbones and promoters are used (Figure 6), creating the possibility of recombination between identical sequences (12) and thus reducing the overall evolutionary stability of the system (36). Furthermore, it is not clear how the cleavage and homing rates will vary as multiplex number is increased substantially. If the gRNA target sites are far away from each other (e.g., >1-5kb), then it is theoretically possible that multiplexing may not be an effective strategy (Supplementary Figure 2A-D), while if they are close together (e.g., <1kb), multiplexing may increase homing effectiveness (Supplementary Figure 2 E-H), although this remains to be demonstrated. Interestingly, a reduction in the homing rate associated with one of the gRNAs may not interfere with our design criteria for population suppression; however, it is important that we have a good quantitative understanding of the underlying molecular dynamics of multiplexed gRNAs in order to make accurate model predictions.

In conclusion, multiplexing gRNAs appears to be a highly effective strategy by which to reduce the effective resistant allele generation rate and hence to enable the elimination of large populations. Due to the exponential decrease in resistant allele generation with increasing multiplex number, only a modest number of gRNAs are needed to achieve population suppression potentially on a continental scale. These approaches need to be tested in relevant organisms to accurately describe their dynamics and to confirm their utility for population suppression strategies. Future studies should address additional sources of resistant alleles, such as *de novo* mutations and naturally-occurring genetic variation. Additional strategies for overcoming resistance should also be explored, for instance, engineering successive gene drive systems each designed to target different essential genes and releasing these one after the other (1). While both this approach and gRNA multiplexing may be effective for overcoming resistance, neither has been demonstrated. Given how quickly this field is advancing, understanding strategies such as these should be of high priority so that the full potential of homing-based population suppression drives can be properly evaluated.

## Materials and methods

### Modeling CRISPR-Cas9 population genetics

To characterize the basic dynamics of the autosomal CRISPR-Cas9-based gene drive system targeting a gene required for female fertility (9), we represent the CRISPR homing construct as a single autosomal allele, “H”, with a corresponding wild-type allele, “h”. We denote homing resistant alleles as “R”. The CRISPR construct creates a bias among gametes of both parental heterozygotes towards gametes having the CRISPR allele. We consider the case where the homing rate is the same among Hh males and females and denote this as *e*. We also consider a resistant allele generation rate that is identical among Hh males and females and denote this as *ρ*. The homing rate, *e*, is the proportion of h gametes in heterozygotes that become H gametes due to the act of homing, and hence the proportion of H gametes arising from heterozygotes of both sexes is equal to (1+*e*)/2. The resistant allele generation rate, *ρ*, is the proportion of h gametes that become R gametes due to errors introduced during the DNA breakage and repair process, and hence the proportion of R gametes arising from heterozygotes of both sexes is equal to *ρ*/2. The remaining gametes arising from heterozygotes are wild-type, h. This leads to an increase in the frequency of the H allele in the population and, since HH females are infertile, there is potential for a population crash to occur under permissive conditions (1, 4). However, since the R allele represents resistance to homing and hence resistance to the spread of the H allele, it has a selective advantage following emergence and is expected to reverse the effects of population suppression. The crosses describing this system are shown in Supplementary Figure 1 and effective resistant allele generation rates for higher multiplex numbers are derived in Supplementary Text S1.

### Modeling An. gambiae population dynamics

Using *An. gambiae* as a case study, we adapt the modeling framework of Deredec *et al.* (5), which itself is based on the population dynamic framework of Hancock and Godfray (37), to describe the spread of the CRISPR and homing-resistant alleles through a discrete, density dependent population with time steps of one day. In this model, the mosquito life cycle is divided into four life stages – egg, larva, pupa and adult (both male and female). The daily, density independent mortality rates for the juvenile stages are assumed to be identical, while the duration of these stages differ. Additional density-dependent mortality occurs at the larval stage, and we use a density-dependent equation of the form, 
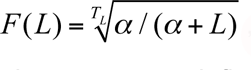
, where *L* is the number of larvae, *T_L_* is the duration of the larval stage *α* is a parameter influencing the strength of density-dependence. Adult males mate throughout their lifetime, while adult females mate only once, soon after that they emerge. Fecundity rates are differ according to genotype, with wild-type females laying *β* eggs per day, females heterozygous for the homing allele laying *β*(1−*s*) eggs per day, HH females being infertile, and females of all other genotypes laying *β* eggs per day. Here, *s* represents the fractional reduction in fertility of females heterozygous for the homing allele. Initial estimates for these and other parameter values are provided in Supplementary Table 1. Equations describing this system are provided in Supplementary Text S1.

We use a stochastic implementation of this model to capture the random effects at low population sizes, for instance when the CRISPR-Cas9 system is causing significant population suppression. We assume that the number of eggs produced per day by females follows a Poisson distribution, the number of eggs having each genotype follows a multinomial distribution, and all survival/death events follow a Bernoulli distribution. Finally, female mate choice follows a binomial distribution with probabilities given by the relative frequency of each male genotype in the population.

### Construct Assembly

Gibson enzymatic assembly (EA) cloning method was used for all cloning (38). To generate plasmid OA-16, components were cloned into the multiple cloning site (MCS) of a commonly used plasmid in the lab for *D. melanogaster* transformation that contains the *white* gene as a marker and an attB-docking site. Specifically, the *Drosophila* ubiquitin promoter (16) was amplified from *D. melanogaster* genomic DNA using primers OA16-1 and OA16-2, and the SV40 3’UTR fragment was amplified from template pMos-3xP3-DsRed-attp (addgene plasmid #52904) using primers OA16-3 and OA16-4. The two RGRs were generated via sequential PCRs using primers OA16-5 and OA16-6 for the first PCR and OA16-5 and OA16-7 for the *white* RGR, and primers OA16-8 and OA16-5 for the first PCR and OA16-9 and OA16-10 for the *yellow* RGR. The construct was assembled in one step: the *D. melanogaster* attB stock plasmid was digested with AscI and XbaI, and the ubiquitin promoter, *white* RGR, *yellow* RGR, and the SV40 3’UTR were cloned in via EA cloning. A list of primer sequences used in the above construct assembly can be found in Supplementary Table 2.

### Fly Culture and Strains

Fly husbandry and crosses were performed under standard conditions at 25°C. Rainbow Transgenics (Camarillo, CA) carried out all of the fly injections. The OA-16 construct was integrated into Bloomington Stock Center (BSC) fly strain 86Fa (BSC #24485: y^1^ M{vasint.Dm}ZH-2A w^*^; M{3xP3-RFP.attP’}ZH-68E), and fly stock BSC#51324 (w[1118]; PBac{y[+mDint2]=vas-Cas9}VK00027) was used as the source of *vasa*-Cas9. For balancing chromosomes, fly stocks BSC#39631 (w[*]; wg[Sp-1]/CyO; P{ry[+t7.2]=neoFRT}82B lsn[SS6]/TM6C, Sb[1]) and BSC#2555 (CyO/sna[Sco]) were used. Homozygous stocks were first generated for 86Fa-OA-16 flies via use of balancer flies. Then, single homozygous female virgins and males were crossed out in triplicate to single male and female virgins, respectively, from the *vasa*-Cas9 line. The offspring (1765 in total) were scored for body color and eye color. The standard error of the mean (SEM) was calculated for each cross type and each phenotype using standard procedures.

### Sequencing to confirm mutations

Genomic DNA was extracted from single mutant flies using Qiagen DNeasy Blood and Tissue Kit (Hilden, Germany), and PCRs were set up using standard protocols to amplify regions of the *white* (primer set OA16-S1/ OA16-S2) and *yellow* (primer set OA16-S3/ OA16-S4) genes that span the cleavage site. Sequencing was performed by Source Bioscience (Nottingham, UK). Primer sequences can be found in Supplementary Table 2.

## Acknowledgements

This work was supported by generous University of California, Riverside (UCR) laboratory start-up funds awarded to O.S.A., a private donation by MaxMind awarded to O.S.A., a US National Institutes of Health (NIH) K22 grant (5K22AI113060-02) awarded to O.S.A., a UC MEXUS grant (CN-15-47) awarded to J.M.M., and a gift from The Parker Foundation to the University of California, San Francisco, Global Health Group Malaria Elimination Initiative.

---

**Supplementary Figure 1.**
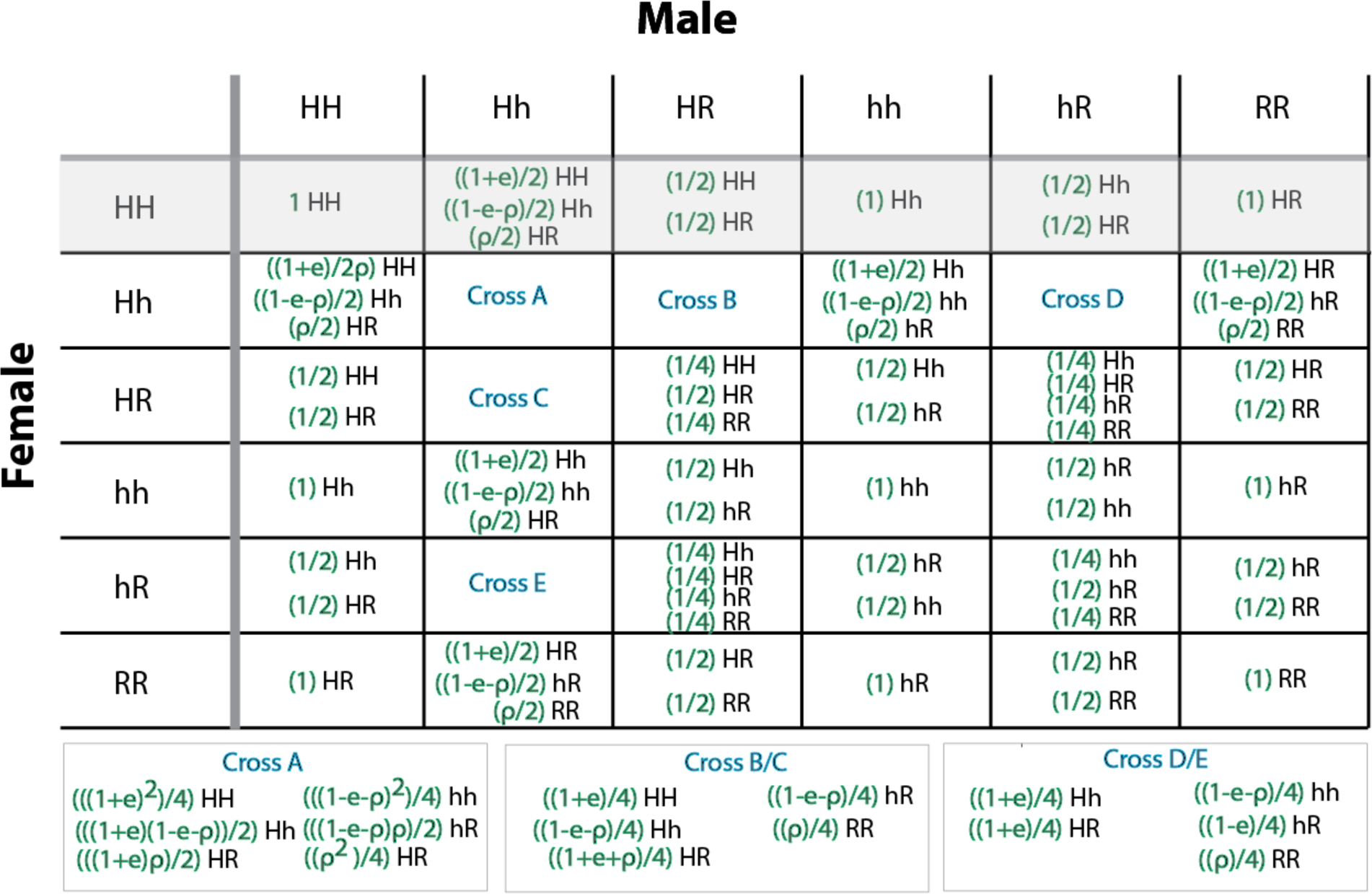
Crosses representing the inheritance pattern of an autosomal CRISPR-Cas9-based homing gene drive system. “H” denotes the CRISPR-Cas9-based homing construct, “h” denotes the corresponding wild-type allele, and “R” denotes a homing resistant allele. Inheritance of the H allele is favored in heterozygous parents as determined by the homing rate, *e*. Homing-resistant alleles may be generated during the process of DNA cleavage and repair at a rate, *ρ*. Crosses involving HH females are shaded out as HH females are rendered infertile by the homing construct. The inheritance pattern of the homing and resistant alleles depicted here is incorporated into the population dynamic model described in the Materials and Methods.

**Supplementary Figure 2.**
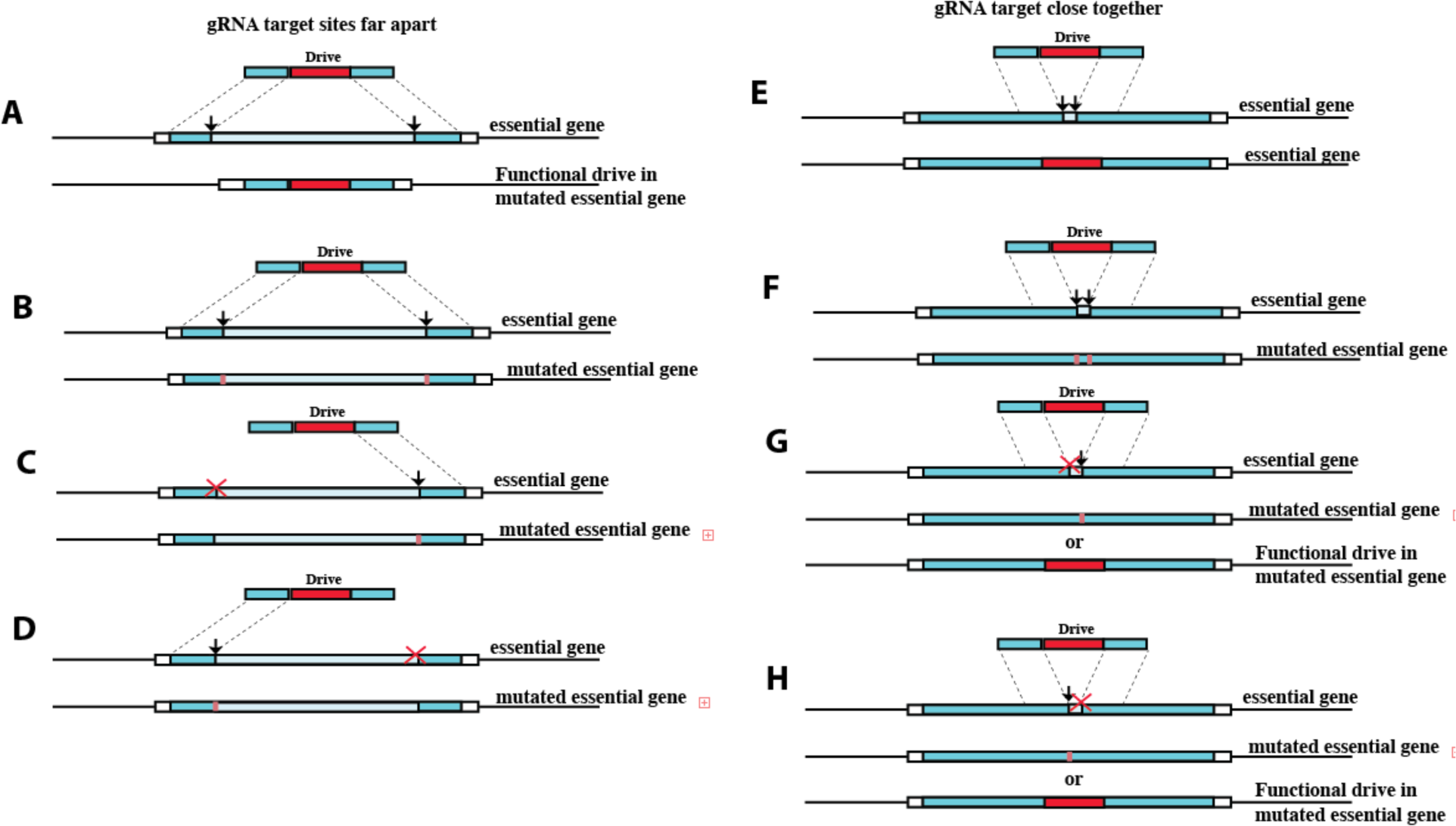
Distance of multiplexed gRNA target sites may affect homing rates. If a suppression based gene drive is designed to target an essential gene at two target sites that are far apart from each other, then four drive-induced possibilities may occur. These include, the drive being successfully copied over to the target allele via HDR following cleavage at both target sites (A), cleavage at both target sites and repair via NHEJ resulting in a mutated drive-resistant essential allele (B), or one of the target sites fails to get cleaved while the other gets cleaved and repaired via NHEJ generating a mutated essential gene (C,D). However, if the drive is designed to target an essential gene at two target sites that are relatively close to each other, but not too close to prevent Cas9 from generating mutations in adjacent target sites, then four different drive-induced possibilities may occur. These include, the drive being successfully copied over to the target allele via HDR following cleavage at both target sites (E), cleavage at both target sites and repair via NHEJ resulting in a mutated drive-resistant essential allele (F), or one of the target sites fails to get cleaved while the other gets cleaved and repaired via NHEJ generating either a mutated essential gene or a functional drive (G,H). Note – for all examples the mutated essential gene may or may not still be functional, however it should be resistant to the endonuclease in the drive due to the mutated gRNA target site.

## Supplementary Text S1

### Model equations for CRISPR-Cas9-based population dynamics in *Anopheles gambiae*

In the main text we describe a stochastic framework for modeling the spread of a CRISPR Cas9-based gene drive system targeting a female fertility gene through a randomly mating population; however equations were left out for brevity. These are included here for completeness.

Using *An. gambiae* as a case study, we adapt the modeling framework of Deredec *et al.* (2011) to describe the spread of the CRISPR and homing-resistant alleles through a discrete, density dependent population with time steps of one day. In this model, the mosquito life cycle is divided into four life stages – egg, larva, pupa and adult (both male and female) – denoted by the subscripts “E”, “L”, “P” and “M”, respectively. The daily, density-independent mortality rates for the juvenile stages are assumed to be identical and are given by *μ*_*E*_ = *μ*_*L*_ = *μ*_*P*_, while the duration of these stages differ and are given by *T_E_*, *T_L_* and *T_P_*. The probability of surviving any of the juvenile stages in a density-independent setting is given by 
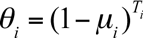, where *i* ∈ {*E*, *L*, *P*}; however additional density-dependent mortality, 1−*F* (*L*), occurs at the larval stage. We use a density-dependent equation of the form, 
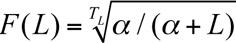, where *α* is a parameter influencing the strength of density-dependence. For adult mosquitoes, the mortality rate is denoted by *μ*_*M*_. Fecundity rates are allowed to differ, with wild-type females laying *β*_*hh*_ = *β* eggs per day, heterozygous females (Hh and HR) laying *β*_*Hh*_ = *β*_*HR*_ = *β*(1−*s*) eggs per day, HH females being infertile (*β*_*HH*_ = 0), and females of other genotypes (hR and RR) laying *β*_*hR*_ = *β*_*RR*_ = *β* eggs per day. Here, *s* represents the fractional reduction in fertility of females heterozygous for the homing allele. Initial estimates for these and other parameter values are provided in Supplementary Table 1.

With this framework in place, the dynamics of the population can be described by equations for the number of larvae and adults belonging to each genotype at time *t*. The number of larvae is needed to determine the strength of density-dependence. Since HH female infertility is irrelevant at the larval stage, we describe the total larval population size at time *t* as,

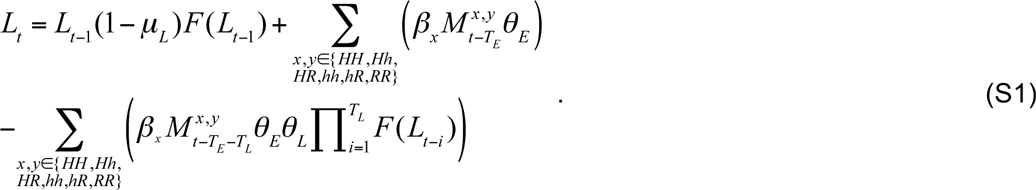

Here, the first term accounts for survival of larvae (denoted at time *t* by *L_t_*) from one day to the next, the second term accounts for newly hatching eggs of any genotype from females of any genotype *x* that have mated with males of any genotype *y* (denoted at time *t* by 
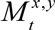), and the third term accounts for transformation of larvae into pupae for juvenile stages resulting from the same crosses.

Adult males and females are treated slightly differently in this framework since it is assumed that female mosquitoes only mate once, while male mosquitoes may mate throughout their lifetime. For example, the number of male adults of genotype HH at time *t* is given by,

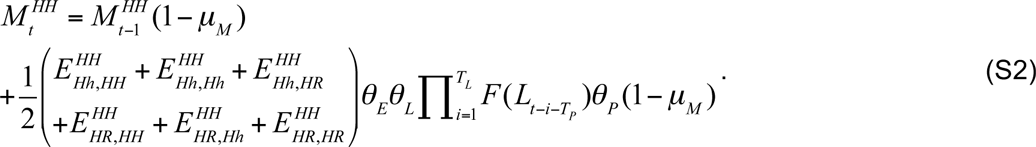

Here, the first term accounts for survival of HH adult males (denoted at time *t* by 
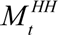) from one day to the next, and the second term accounts for transformation of HH pupae into adult males, where these pupae result from crosses between Hh and HR females with HH, Hh and HR males. The number of eggs of genotype *x* produced by adult females of genotype *y* that have mated with a male of genotype *z* is given by 
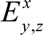. These quantities are time-dependent and the product of the fecundity of the female genotype, *β*_*y*_, the number of females having the given mated genotype, *M^y,z^*, and the proportion of offspring of this mated genotype having the genotype *z* (depicted in the crosses shown in Supplementary Figure 1). The numbers of HH eggs from each cross in Equation S2 are given by the following equations:

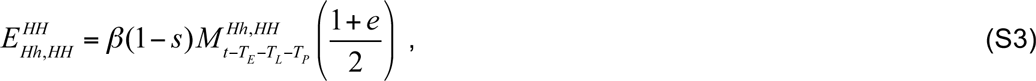

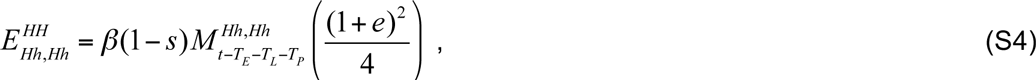

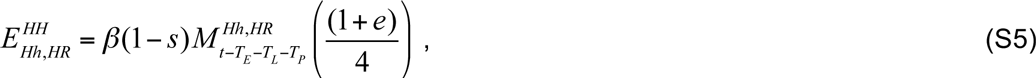

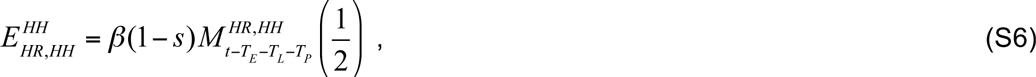

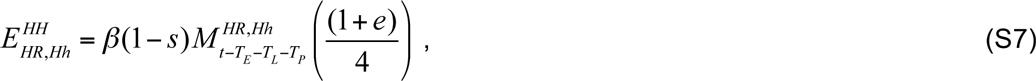

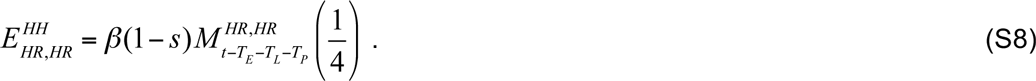

Crosses involving HH females are not included here, as these females are rendered infertile by the CRISPR construct.

Females, on the other hand, are assumed to mate only once and on the same day that they emerge. They can therefore be described by both their genotype and the genotype of the male with whom they mated. For example, the number of female adults at time *t* of genotype HH that have mated with hh males is given by,

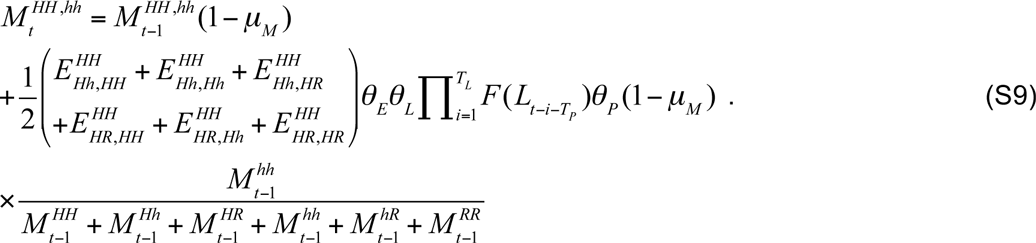

Here, the first term accounts for survival of HH adult females that have mated with hh males (denoted at time *t* by 
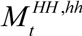) from one day to the next, and the second term accounts for transformation of HH pupae into adult females, where these pupae result from crosses between Hh and HR females with HH, Hh and HR males. This term is multiplied by the fraction of the adult male population having the genotype hh. Equations for all other adult genotypes are treated analogously as follows.

Equation S2 describes the number of adult males of genotype HH over time. There are five other male genotypes – Hh, HR, hh, hR and RR – denoted by the variables 
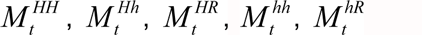 and 
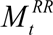 respectively, and described by the following equations:

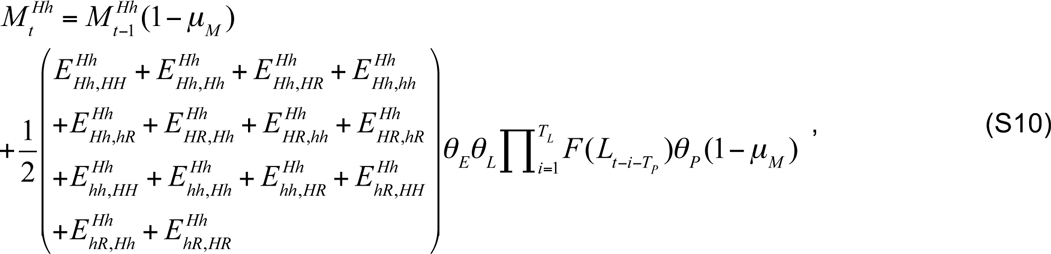

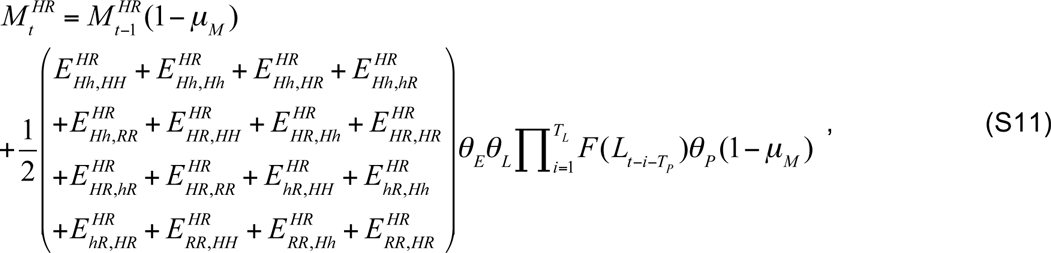

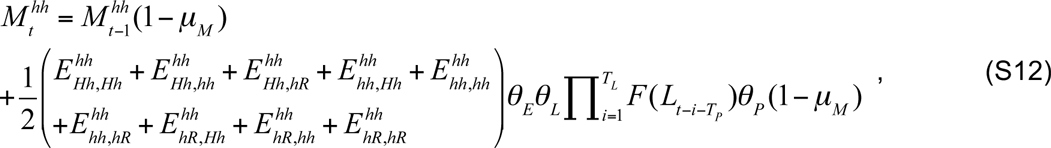

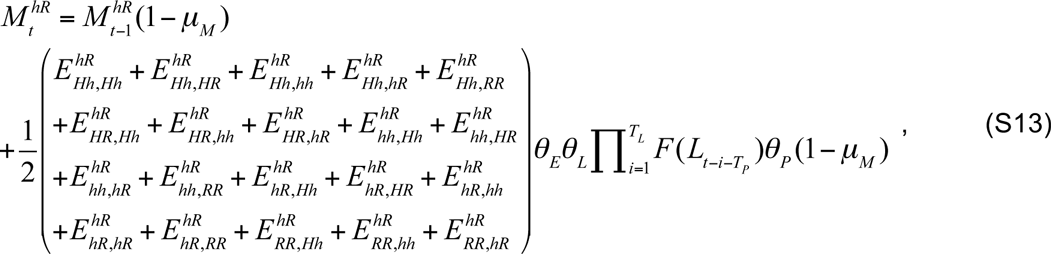

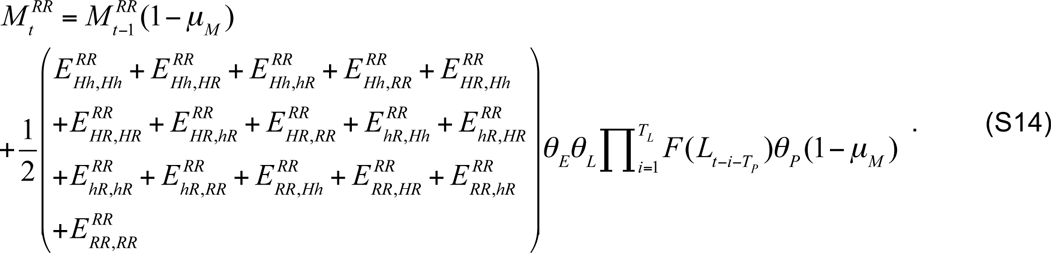

Here, the first term accounts for adult survival for each genotype from one day to the next, and the second term accounts for transformation of pupae into adult males for each genotype, where pupae result from the crosses depicted in Supplementary Figure 1. The number of eggs of genotype *x* produced by adult females of genotype *y* that have mated with a male of genotype *z* is given by 
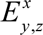. These quantities are the product of the fecundity of the female genotype, *β*_*y*_, the number of females having the given mated genotype, *M*^*y,z*^, and the proportion of offspring of this mated genotype having the genotype *z*. The numbers of eggs of all genotypes for each of the crosses depicted in Supplementary Figure 1 are given by the following equations:

Eggs produced by *Hh* females:

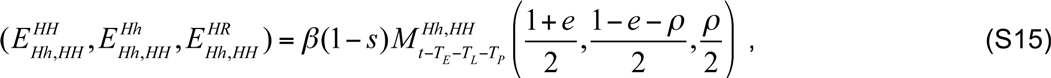

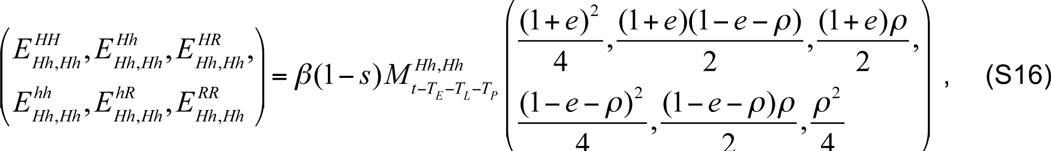

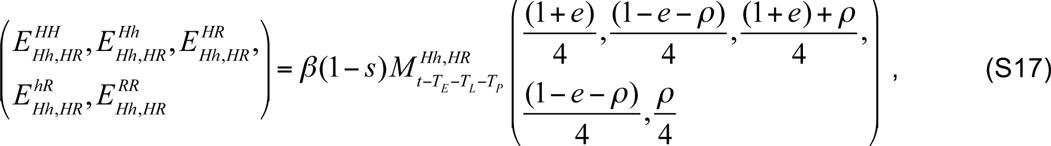

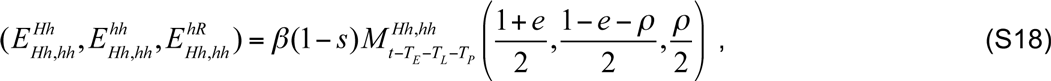

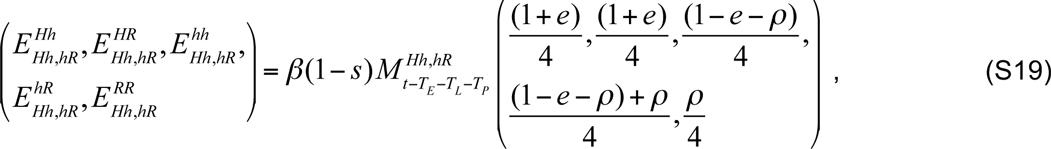

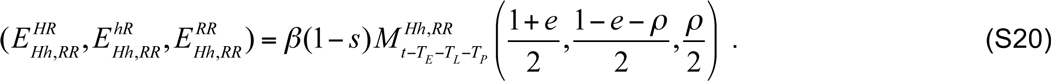

Eggs produced by *HR* females:

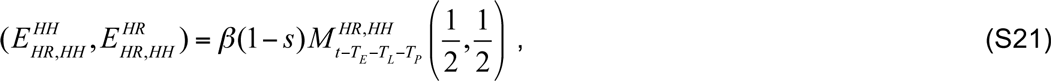

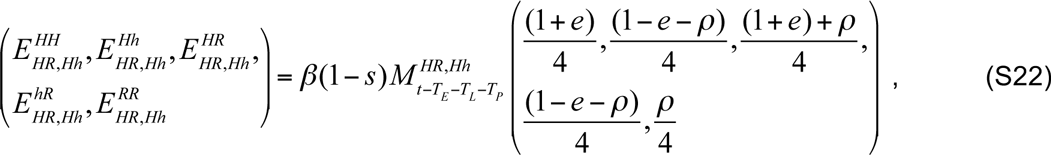

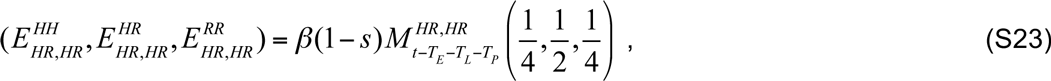

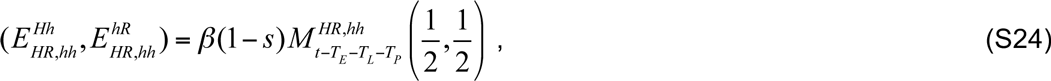

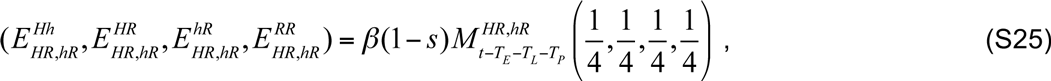

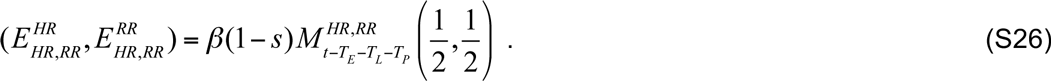

Eggs produced by *hh* females:

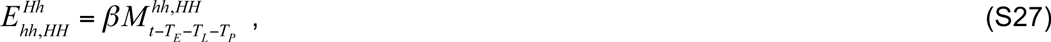

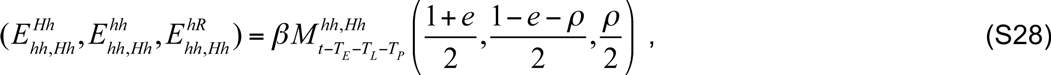

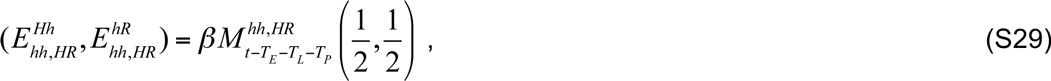

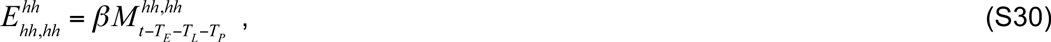

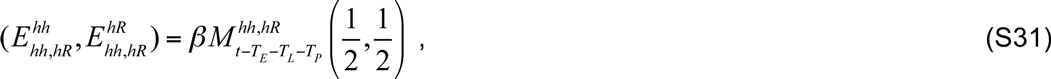

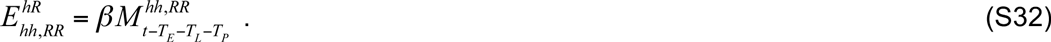

Eggs produced by *hR* females:

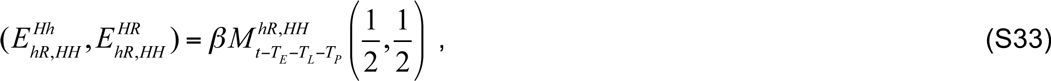

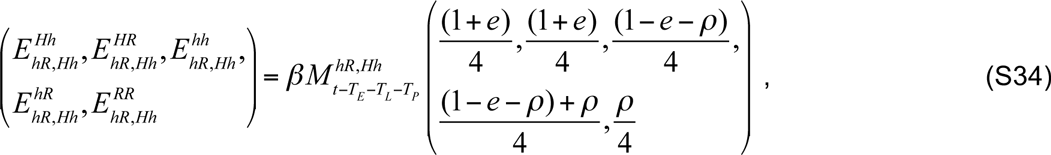

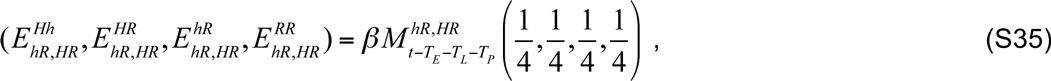

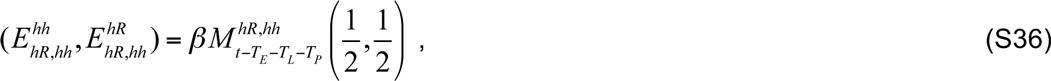

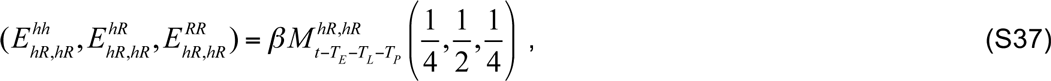

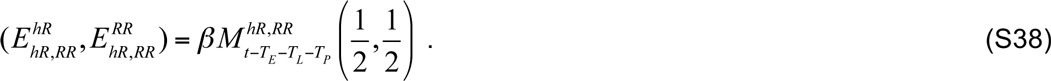

Eggs produced by *RR* females:

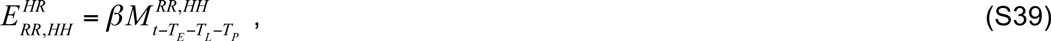

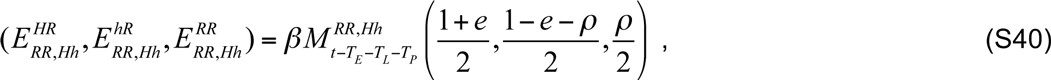

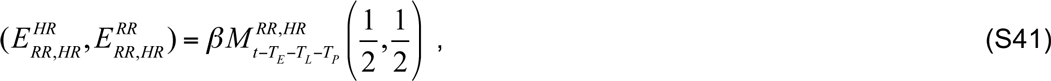

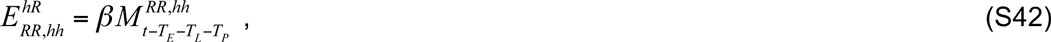

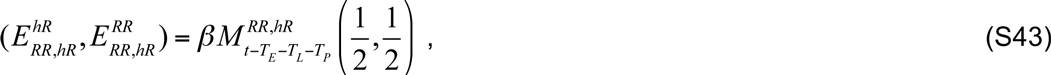

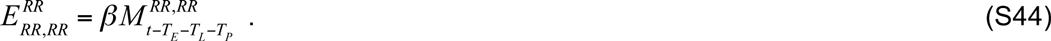

As mentioned earlier, crosses involving HH females are not included here, as these females are rendered infertile by the CRISPR construct.

Females are assumed to mate only once and on the same day that they emerge so can therefore be described by both their genotype and the genotype of the male with whom they mated. Equation S9 describes the number of female adults of genotype HH that have mated with hh males over time. The other mated female genotypes are described by the following equations:

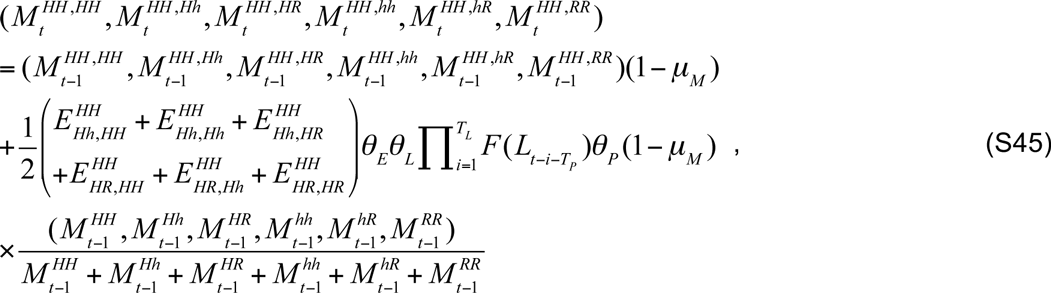

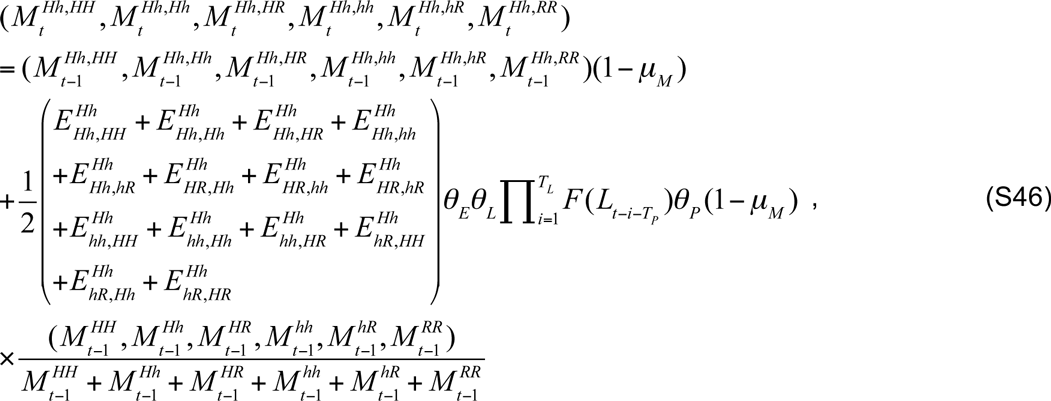

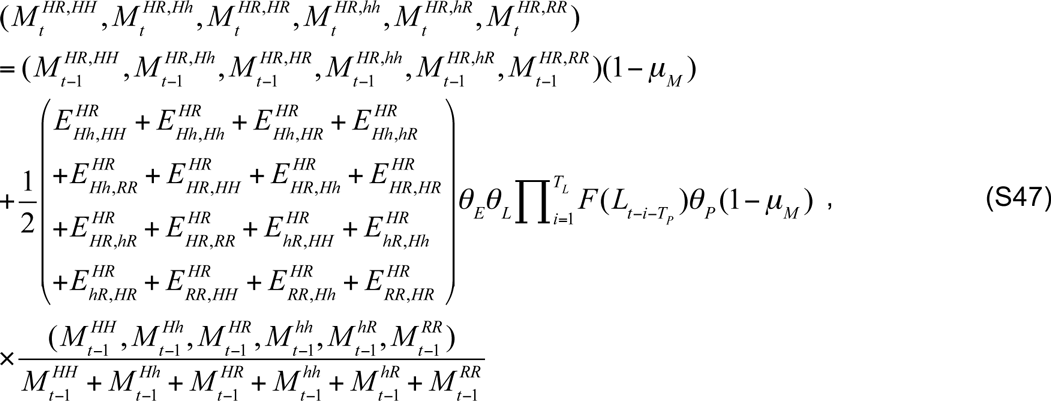

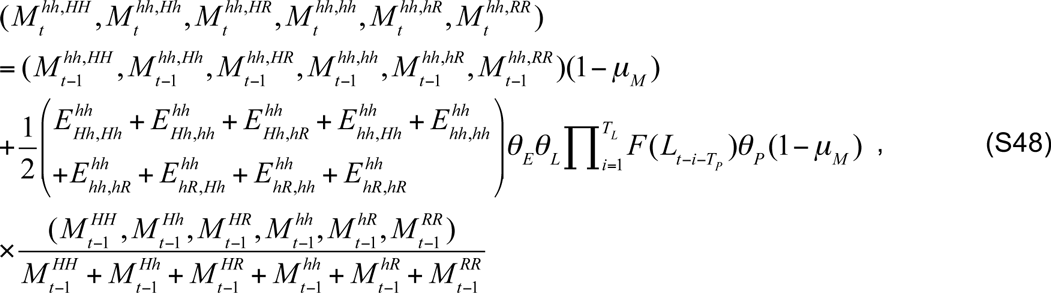

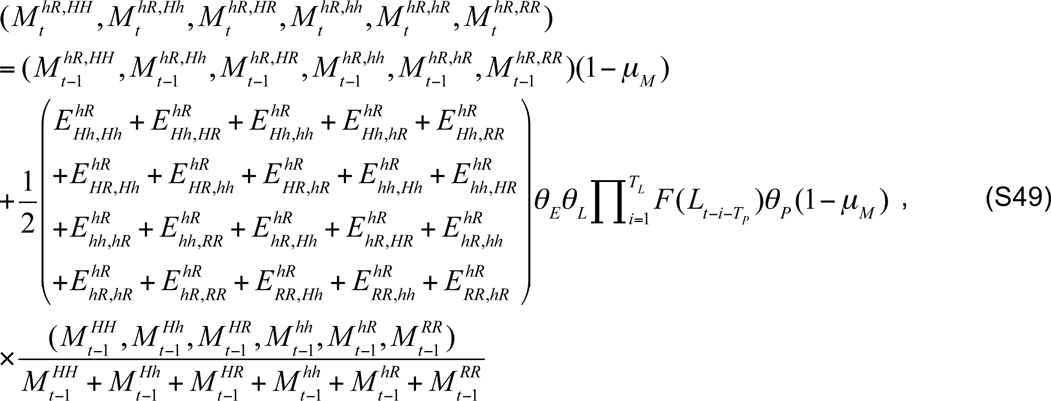

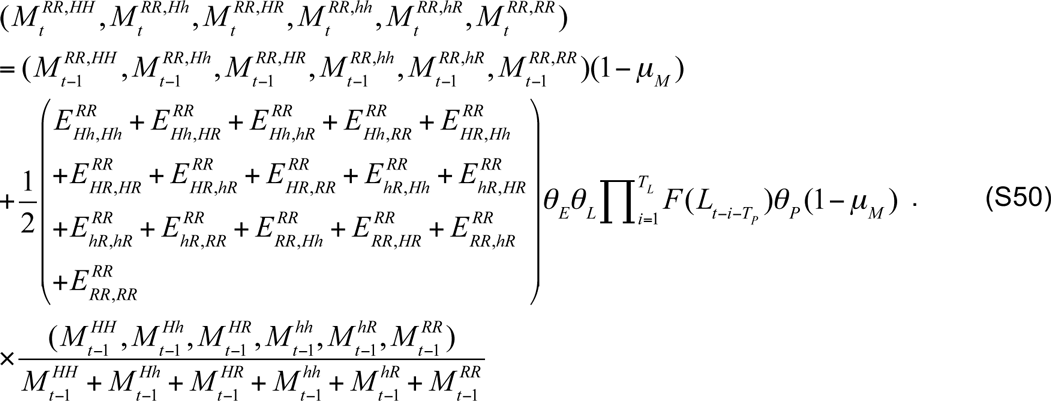

For each of these equations, the first term accounts for survival of adult females having the given mated genotype from one day to the next, and the second term accounts for transformation of pupae of the given female genotype into adults. The second term is then multiplied by the fraction of the adult male population having either genotype HH, Hh, HR, hh, hR or RR, depending on the female mated genotype.

Using these equations, we can derive several basic properties of the population, such as the non-zero equilibrium densities of larvae and adults, and the basic reproductive number – i.e. the average number of female offspring produced by a single female that survive to adulthood at low population densities – in the absence of genetic control. The basic reproductive number is equal to the rate of female egg production multiplied by the life expectancy of an adult mosquito multiplied by the proportion of eggs that will survive through all of the juvenile life stages in the absence of density-dependence. This is given by,

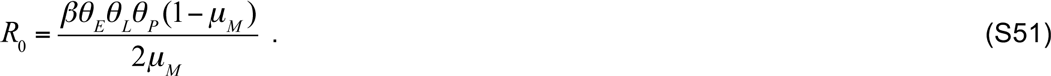

The equilibrium population densities can then be calculated by setting the population densities to be equal across generations in Equations S1, S12, S30 and S48. This leads to the following non-zero equilibria:

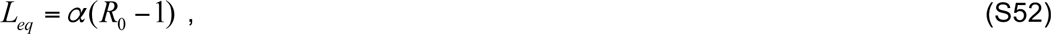

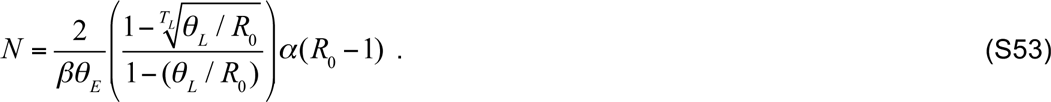

Here, *L_eq_* and *N* represent the total population equilibria (i.e. 
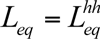 and 
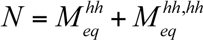. These formulations guide the parameter choices, taken from Deredec *et al.* (2011), as shown in Table S1:

**Table S1:** Parameter values for stochastic, discrete-time model.

| Symbol: | Parameter: | Value: | References: |
| --- | --- | --- | --- |
| Primary parameters: |  |  |  |
| $\beta$ | Egg production per wild-type female | 32 /day | Depinay *et al.* (2004) |
| $T_E$ | Duration of egg stage | 1 day | Depinay *et al.* (2004) |
| $T_L$ | Duration of larval stage | 14 days | Depinay *et al.* (2004) |
| $T_P$ | Duration of pupal stage | 1 day | Depinay *et al.* (2004) |
| $\mu_E = \mu_L = \mu_P$ | Death rate of juvenile stages | 0.168 /day | Molineaux & Gramiccia (1980),<br>Depinay *et al.* (2004) |
| $\mu_M$ | Death rate of adult stage | 0.123 /day | Molineaux & Gramiccia (1980) |

| Variable parameters: |  |  |  |
| --- | --- | --- | --- |
| $e$ | Homing rate | [0.98, 0.9999] | Hammond *et al.* (2016) |
| $\rho$ | Resistant allele generation rate | $[10^{-7}, 10^{-2}]$ | Hammond *et al.* (2016) |
| $s$ | Fertility cost to Hh females | 0, 0.907 | Hammond *et al.* (2016) |
| $N$ | Equilibrium adult mosquito population size (male and female) | $[10^3, 10^6]$ | |

### Modeling gRNA multiplexing

The results depicted in Figure 3 in the main text suggest that the probability of population elimination is independent of the homing rate for *e* > 98%, but is highly dependent on the resistant allele generation rate, *ρ*. We propose gRNA multiplexing as a method to reduce the effective resistant allele generation rate and, here, describe how the effective resistant allele generation rate varies with multiplex number.

#### Multiplex number of two

For the example of two multiplexed gRNAs, there are two sites within a composite allele at which the gRNAs cleave, both of which may either have the homing construct, H, be resistant to homing, R, or be wild-type, h. We denote the composite allele as {*xy*}, where *x* denotes the first site in the composite allele, *y* denotes the second side in the composite allele, and *x*, *y* ∈ {H, R, h}.

As multiplexing provides multiple opportunities for homing to occur, we consider a composite allele to have the homing phenotype (the ability to cleave and home into the homologous chromosome at multiple target sites) if at least one of its sites has a functional copy of the homing allele (i.e. the composite alleles {HH}, {Hh} and {HR}). We consider a composite allele to have the homing-resistant phenotype if all of its sites have a homing-resistant allele (i.e. the composite allele {RR}). All other composite alleles are considered to have the wild-type phenotype, i.e. they don’t have the homing phenotype but are still potentially receptive to a homing event. For the two-gRNA system, the effective resistant allele generation rate, *ρ*_*m*=2_, is then given by,

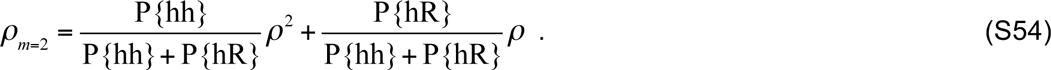

Here, P{hh} and P{hR} represent the proportion of composite alleles that are {hh} and {hR}, respectively, *ρ*^2^ is the probability of generating a {RR} composite allele from a {hh} composite allele when the composite allele on the opposite chromosome has at least one functional copy of the homing allele, and *ρ* is the probability of generating an {RR} allele from an {hR} allele under the same circumstances.

The relative proportion of {hh} and {hR} composite alleles can be estimated by considering the flux of composite alleles as they become associated with composite alleles on the opposite chromosome having the homing phenotype. For the two-gRNA system, the flux of {hR} composite alleles is given by,

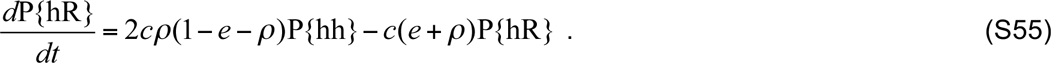

Here, *c* represents the rate at which a given chromosome becomes associated with an opposite chromosome having the homing phenotype. The rate of change of the proportion of chromosomes having the {hR} composite allele is equal to the rate at which {hR} composite alleles are generated from {hh} alleles (i.e. through resistant allele generation at one site and wild-type allele maintenance at the other site) subtracting the rate at which {hR} composite alleles are lost through either homing or resistant allele generation at the remaining wild-type site.

Allele frequencies are constantly in flux as a gene drive system spreads into a population; however, if we assume, to a first approximation, that equilibrium is maintained between {hh} and {hR} composite alleles for a given prevalence of composite alleles having the homing phenotype, then the equilibrium solution to Equation S55 suggests the following ratio of {hR} to {hh} composite alleles:

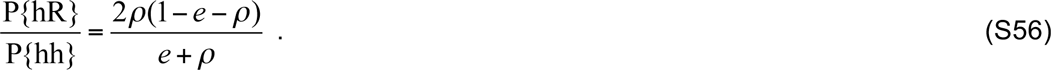

Substituting this ratio into Equation S54, the effective resistant allele generation rate for two multiplexed gRNAs, *ρ*_*m*=2_, is given by,

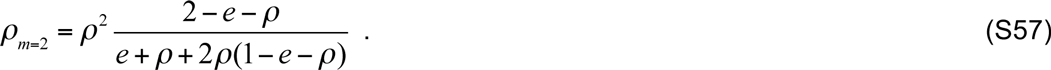

Substituting our homing efficiency of 98% and resistant allele generation rate of 1% into Equation S57, we see that the effective resistant allele generation rate for a multiplex number of two is equal to *ρ*^2^ multiplied by a fraction very close to 1 (1.02). I.e. for the circumstances being studied,

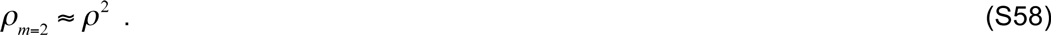

This is a consequence of most resistant allele generation occurring directly from {hh} composite alleles, since {hR} composite alleles are rarely generated and are frequently converted to alleles having the homing phenotype once they have been formed.

#### Multiplex number of three

To check whether the same approximation holds for three multiplexed gRNAs, i.e. that most resistant allele generation occurs directly from {hhh} composite alleles, we calculate the effective resistant allele generation rate using the same framework as described above. We denote the composite allele in this case as {*xyz*}, where *x*, *y*, *z* ∈ {H, R, h}. As multiplexing provides multiple opportunities for homing to occur, a composite allele is considered to have the homing phenotype if at least on of its sites has a functional copy of the homing allele, a composite allele is considered to have the homing-resistant phenotype if all of its sites have a homing-resistant allele, and all other composite alleles have the wild-type phenotype.

For the three-gRNA system, the effective resistant allele generation rate, *ρ*_*m*=3_, is given by,

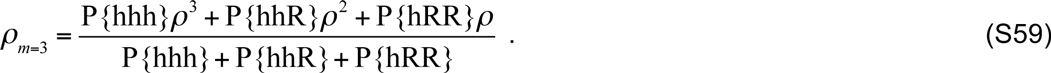

Here, P{hhh}, P{hhR} and P{hRR} represent the proportion of composite alleles that are {hhh}, {hhR} and {hRR}, respectively, and *ρ*^3^, *ρ*^2^ and *ρ* are the probabilities of generating an {RRR} composite allele from an {hhh}, {hhR} and {hRR} composite allele, respectively, when the corresponding allele has at least one functional copy of the homing allele.

As for the two-gRNA case, the relative proportion of {hhh}, {hhR} and {hRR} composite alleles can be estimated by considering the relative flux of allele genotypes as they become associated with corresponding composite alleles having the homing phenotype. For the three-gRNA system, the flux of the {hhR} composite alleles is given by,

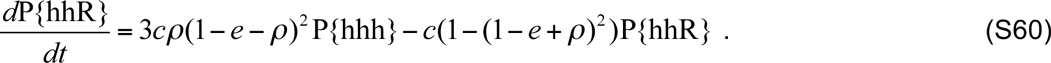

Here, the rate of change of the proportion of chromosomes having the {hhR} composite allele is equal to the rate at which {hhR} composite alleles are generated from {hhh} composite alleles (i.e. through resistant allele formation at one site and wild-type allele maintenance at the other two sites) subtracting the rate at which {hhR} composite alleles are lost through either homing or resistant allele generation at the remaining two wild-type sites. Assuming, to a first approximation, that equilibrium is maintained between {hhh} and {hhR} composite alleles for a given prevalence of composite alleles having the homing phenotype, then the equilibrium solution to Equation S60 suggests the following ratio of {hhR} to {hhh} composite alleles:

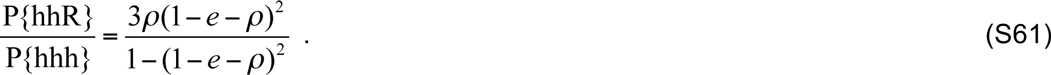

Similarly, the flux of {hRR} composite alleles is given by,

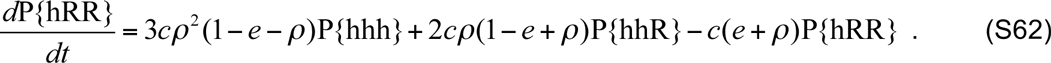

Here, the rate of change of the proportion of chromosomes having the {hRR} composite allele is equal to the rate at which {hRR} composite alleles are generated from {hhh} composite alleles (i.e. through resistant allele formation at two sites and wild-type allele maintenance at the other site) added to the rate at which {hRR} composite alleles are generated from {hhR} composite alleles (i.e. through resistant allele formation at one site and wild-type allele maintenance at the other site) subtracting the rate at which {hRR} composite alleles are lost through either homing or resistant allele generation at the remaining wild-type site. Substituting Equation S61 into Equation S62 and assuming, to a first approximation, that equilibrium is maintained between {hhh} and {hRR} alleles for a given prevalence of composite alleles having the homing phenotype, then the equilibrium solution to Equation S62 suggests the following ratio of {hRR} to {hhh} composite alleles:

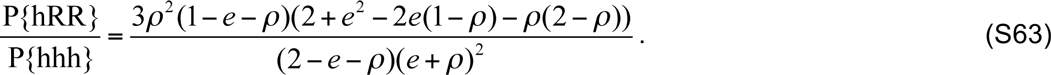

Substituting the ratios in Equations S61 and S63 into Equation S59, the effective resistant allele generation rate for three multiplexed gRNAs, *ρ*_*m*=3_, is given by,

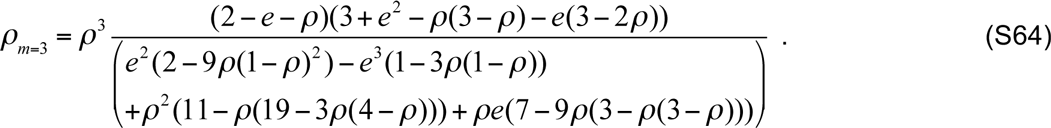

Substituting our homing efficiency of 98% and resistant allele generation rate of 1% into Equation S64, we see that the effective resistant allele generation rate for a multiplex number of three is equal to *ρ*^3^ multiplied by a fraction very close to 1 (1.03). I.e. for the circumstances being studied,

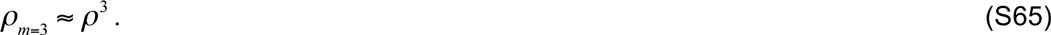

This is a consequence of most resistant allele generation occurring directly from {hhh} composite alleles, since {hhR} and {hRR} composite alleles are rarely generated and are frequently converted to composite alleles having the homing phenotype once they have been formed. We have reason to believe this trend will continue for higher multiplex numbers for the parameter ranges we are exploring here. Therefore, for the purposes of this paper, we will approximate the effective resistant allele generation rate for a multiplex number of *m* as *ρ_m_*.

#### Multiplex number of two (reduced homing rate for second gRNA)

In the above calculations, we have ignored the reduced cleavage rate observed for the RGR multiplexing approach in *D. melanogaster* for the second gRNA. If we assume a fractional reduction in cleavage rate, *f*, and that this will reduce both the homing and resistant allele generation rates by the same amount for the second gRNA, then we can derive how this effects the effective resistant allele generation rate by modifying the previous analysis for two multiplexed gRNAs. We will now consider ordered composite alleles, where {hR} represents a composite allele with a wild-type allele at the first site and a homing-resistant allele at the second site, and {Rh} represents a composite allele with a homing-resistant allele at the first site and a wild-type allele at the second site. For the two-gRNA system, the effective resistant allele generation rate, *ρ*_*m*=2_, is now given by,

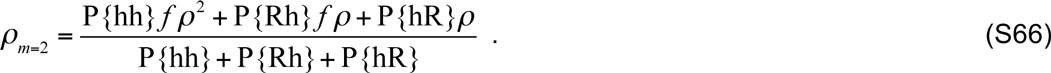

Here, *fρ*^2^ is the probability of generating a {RR} composite allele from a {hh} composite allele when the composite allele on the opposite chromosome has at least one functional copy of the homing allele, *ρ* is the probability of generating a {RR} composite allele from a {hR} composite allele under the same circumstances, and *fρ* is the probability of generating a {RR} composite allele from a {Rh} composite allele, where the reduction is due to the reduced cleavage rate at the second site.

The relative proportions of {hh}, {Rh} and {hR} composite alleles can be estimated by considering the flux of composite alleles as they become associated with composite alleles on the opposite chromosome having the homing phenotype. For the two-gRNA system with reduced cleavage rate at site two, these are given by,

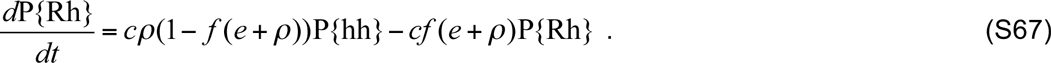

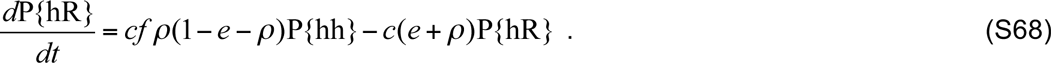

Here, {Rh} composite alleles are generated at a faster rate due to the higher likelihood that the second site will remain wild-type, and {hR} composite alleles are generated at a slower rate due to the smaller likelihood that cleavage and hence resistant alleles will occur at the second site. {Rh} composite alleles are also lost at a slower rate due to homing and resistant allele generation occurring at a slower rate at the second site.

If we assume, to a first approximation, that equilibrium is maintained between {hh}, {Rh} and {hR} composite alleles for a given prevalence of composite alleles having the homing phenotype, then the equilibrium solution to Equations S67-S68 suggests the following ratio of {Rh} and {hR} to {hh} composite alleles:

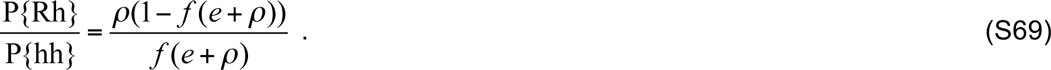

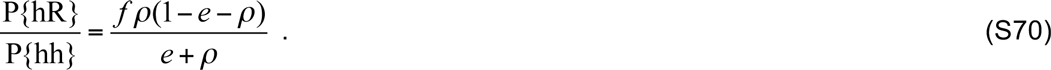

Substituting these ratios into Equation S66, the effective resistant allele generation rate for two multiplexed gRNAs, *ρ*_*m*=2_, with reduced cleavage at site two is given by,

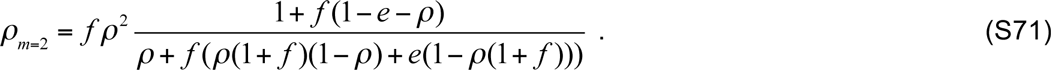

Substituting our homing efficiency of 98% and baseline *ρ* value of 1% into Equation S71, we see that, for a reduced cleavage rate of 75% at site two, the effective resistant allele generation rate is equal to *ρ*^2^ multiplied by 1.014, and for a reduced cleavage rate of 50% at site two, the effective resistant allele generation rate is *ρ*^2^ multiplied by a 1.005. In fact, even for a drastically reduced cleavage rate of 25% at site two, the effective resistant allele generation rate is still approximately *ρ*^2^ (*ρ*^2^ multiplied by a 0.983). Thus, interestingly, a reduction in cleavage rate at site two doesn’t significantly alter the effective resistant allele generation rate, and in fact slightly reduces the rate as compared to that without a reduced cleavage rate at the second site (which was *ρ*^2^ multiplied by 1.020).

**Table S2:**
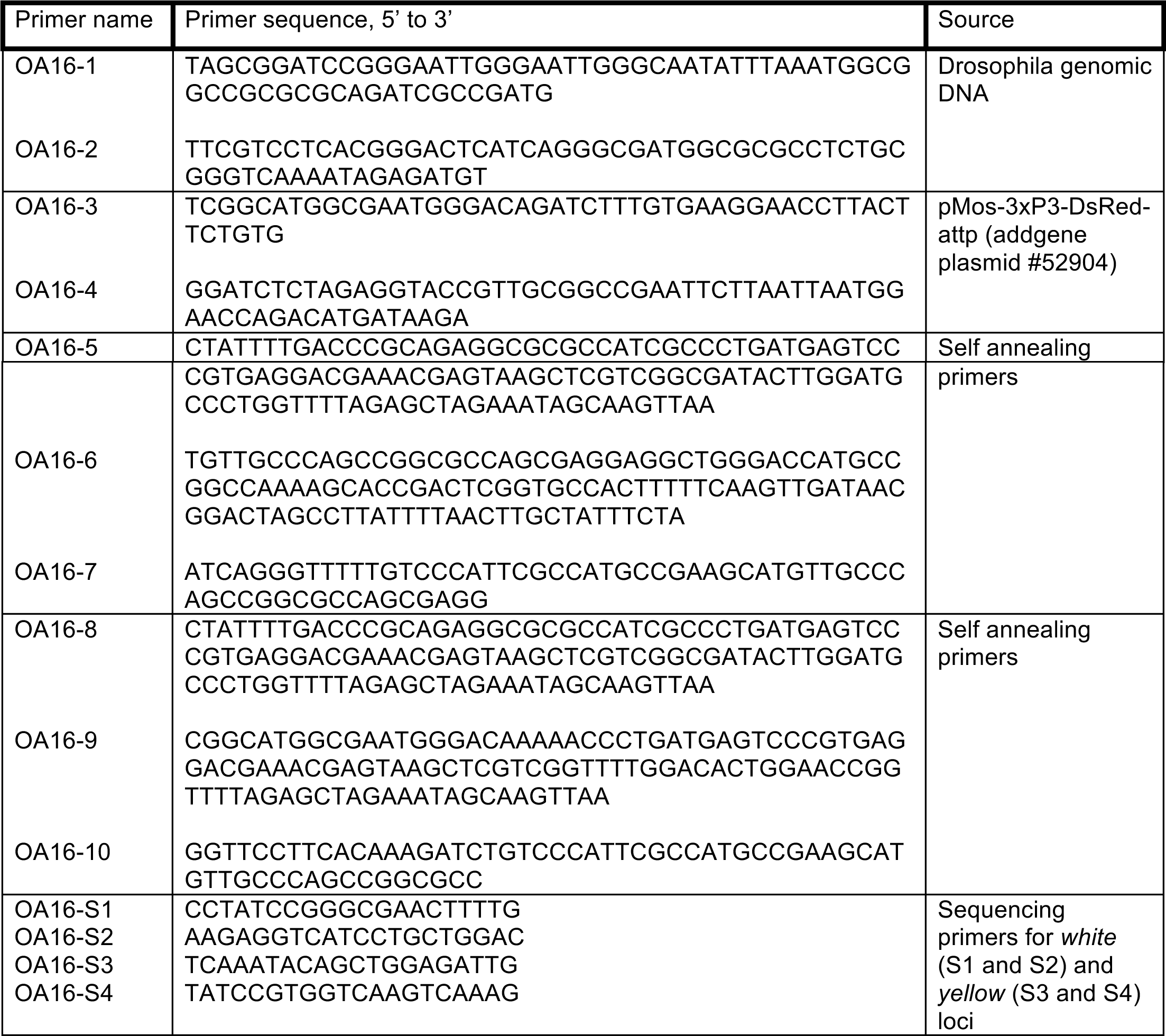
Primer sequences Sequences

